# Enhanced skeletal muscle ribosome biogenesis, yet attenuated mTORC1 and ribosome biogenesis-related signalling, following short-term concurrent versus single-mode resistance training

**DOI:** 10.1101/115212

**Authors:** Jackson J. Fyfe, David J. Bishop, Jonathan D. Bartlett, Erik D. Hanson, Mitchell J. Anderson, Andrew P. Garnham, Nigel K. Stepto

**Author notes:** **Address for correspondence:** Jackson J. Fyfe, PhD, School of Exercise and Nutrition Sciences, Deakin University, 221 Burwood Hwy, Burwood VIC 3125.

## Abstract

Combining endurance training with resistance training (RT) may attenuate skeletal muscle hypertrophic adaptation versus RT alone; however, the underlying mechanisms are unclear. We investigated changes in markers of ribosome biogenesis, a process linked with skeletal muscle hypertrophy, following concurrent training versus RT alone. Twenty-three males underwent eight weeks of RT, either performed alone (RT group, *n* = 8), or combined with either high-intensity interval training (HIT+RT group, *n* = 8), or moderate-intensity continuous training (MICT+RT group, *n* = 7). Muscle samples (*vastus lateralis*) were obtained before training, and immediately before, 1 h and 3 h after the final training session. Training-induced changes in basal expression of the 45S ribosomal RNA (rRNA) precursor (45S pre-rRNA), and 5.8S and 28S mature rRNAs, were greater with concurrent training versus RT. However, during the final training session, RT further increased both mTORC1 (p70S6K1 and rps6 phosphorylation) and 45S pre-rRNA transcription-related signalling (TIF-1A and UBF phosphorylation) versus concurrent training. These data suggest that when performed in a training-accustomed state, RT induces further increases mTORC1 and ribosome biogenesis related signalling in human skeletal muscle versus concurrent training; however, changes in ribosome biogenesis markers were more favourable following a period of short-term concurrent training versus RT performed alone.

## 2. Introduction

Simultaneously incorporating both resistance and endurance training into a periodised training program, termed concurrent training ^1^, can attenuate resistance training adaptations such as muscle hypertrophy, compared with resistance training performed alone ^2-4^. This effect is potentially mediated by an altered balance between post-exercise skeletal muscle protein synthesis (MPS) and breakdown, subsequently attenuating lean mass accretion. The mechanistic target of rapamycin complex 1 (mTORC1) is a key mediator of load-induced increases in MPS and subsequently muscle hypertrophy ^5,6^. The activity of mTORC1 is antagonised by activation of the 5’ adenosine monophosphate-activated protein kinase (AMPK), which acts to restore perturbations in cellular energy balance by inhibiting anabolic cellular processes and stimulating catabolism ^7^. For example, in rodent skeletal muscle, low frequency electrical stimulation mimicking endurance exercise-like contractions promotes AMPK activation and inhibition of mTORC1 signalling ^8^.

Subsequent work in humans ^9-18^ has focused on the hypothesis that attenuated muscle hypertrophy with concurrent training ^2,4,19^ may be explained by AMPK-mediated inhibition of the mTORC1 pathway. Several studies, however, have demonstrated that single sessions of concurrent exercise do not compromise either mTORC1 signalling or rates of MPS ^9,10,16-18^, and may even potentiate these responses ^14^, compared with resistance exercise performed alone. However, a limitation of these studies is that most have examined these responses in either untrained individuals ^16-18^ or those who are relatively unaccustomed to the exercise protocol ^14,20^. Given short-term training increases the mode-specificity of post-exercise molecular responses ^21,22^, examining perturbations to molecular signalling and gene expression in relatively training-unaccustomed individuals may confound any insight into the potential molecular mechanisms responsible for interference following concurrent training ^23^.

Transient changes in translational efficiency (i.e., rates of protein synthesis per ribosome) after single sessions of concurrent exercise, as indexed by skeletal muscle mTORC1 signalling or rates of MPS, in relatively training-unaccustomed individuals therefore do not appear to explain interference to muscle hypertrophy following longer-term concurrent training. However, rates of cellular protein synthesis are determined not only by transient changes in translational efficiency, but also by cellular translational capacity (i.e., amount of translational machinery per unit of tissue, including ribosomal content) ^24^. Ribosomes are supramolecular ribonucleoprotein complexes functioning at the heart of the translational machinery to convert mRNA transcripts into protein ^24^, and ribosomal content dictates the upper limit of cellular protein synthesis ^25^. Early rises in protein synthesis in response to anabolic stimuli (e.g., a single bout of resistance exercise) are generally thought to be mediated by transient activation of existing translational machinery, whereas prolonged anabolic stimuli (e.g., weeks to months of RE training) induces an increase in total translational capacity via ribosome biogenesis ^24^.

Ribosome biogenesis is a complex, well-orchestrated process involving transcription of the polycistrionic 45S rRNA (ribosomal RNA) precursor (45S pre-rRNA), processing of the 45S pre-rRNA into several smaller rRNAs (18S, 5.8S and 28S rRNAs), assembly of these rRNAs and other ribosomal proteins into ribosomal subunits (40S and 60S), and nuclear export of these ribosomal subunits into the cytoplasm ^24,26^. The synthesis of the key components of the ribosomal subunits is achieved via the coordinated actions of three RNA polymerases (RNA Pol-I, -II, and -III). The RNA Pol-I is responsible for the transcription of the 45S pre-rRNA in the nucleolus, which is considered the rate-limiting step in ribosome biogenesis ^27^. The 45S pre-rRNA is subsequently cleaved into the 18S, 5.8S and 28S rRNAs, which undergo post transcriptional modifications via interactions with small nuclear ribonucleoproteins and several protein processing factors. The RNA Pol-II is responsible for the transcription of ribosomal protein-encoding genes, whereas RNA Pol-III mediates the nucleoplasmic transcription of 5S rRNA and tRNAs (transfer RNAs) ^26^.

As well as controlling translational efficiency, the mTORC1 is a key mediator of ribosome biogenesis by regulating transcription factors for genes encoding RNA Pol-I (see Figure 1) and -III ^25^. The transcription of rDNA by RNA Pol-I requires the transcription factor SL-1 (selectivity factor-1), a component of which is TIF-1A (transcription initiation factor 1A; also known as RRN5), as well as other regulatory factors including POLR1B (polymerase [RNA] 1 polypeptide B). Inhibition of mTORC1 by rapamycin inactivates TIF-1A, which impairs the transcription of the 45S pre-rRNA by RNA Pol-I ^28^. Inhibition of mTORC1 also inactivates UBF (upstream binding factor) ^29^, a transcription factor also associated with SL-1, while the key mTORC1 substrate p70S6K1 promotes UBF activation and RNA Pol-I-mediated rDNA transcription ^29^. As well as regulation by mTORC1 signalling, the cyclins (including cyclin D1) and cyclin-dependent kinases (CDKs) can also regulate UBF via phosphorylation on Ser388 and Ser484, which are required for UBF activity ^30,31^. In addition to regulation of RNA Pol-1, mTORC1 also associates with a number of RNA Pol-III genes that synthesise 5S rRNA and tRNA ^32^.

**Figure 1.**
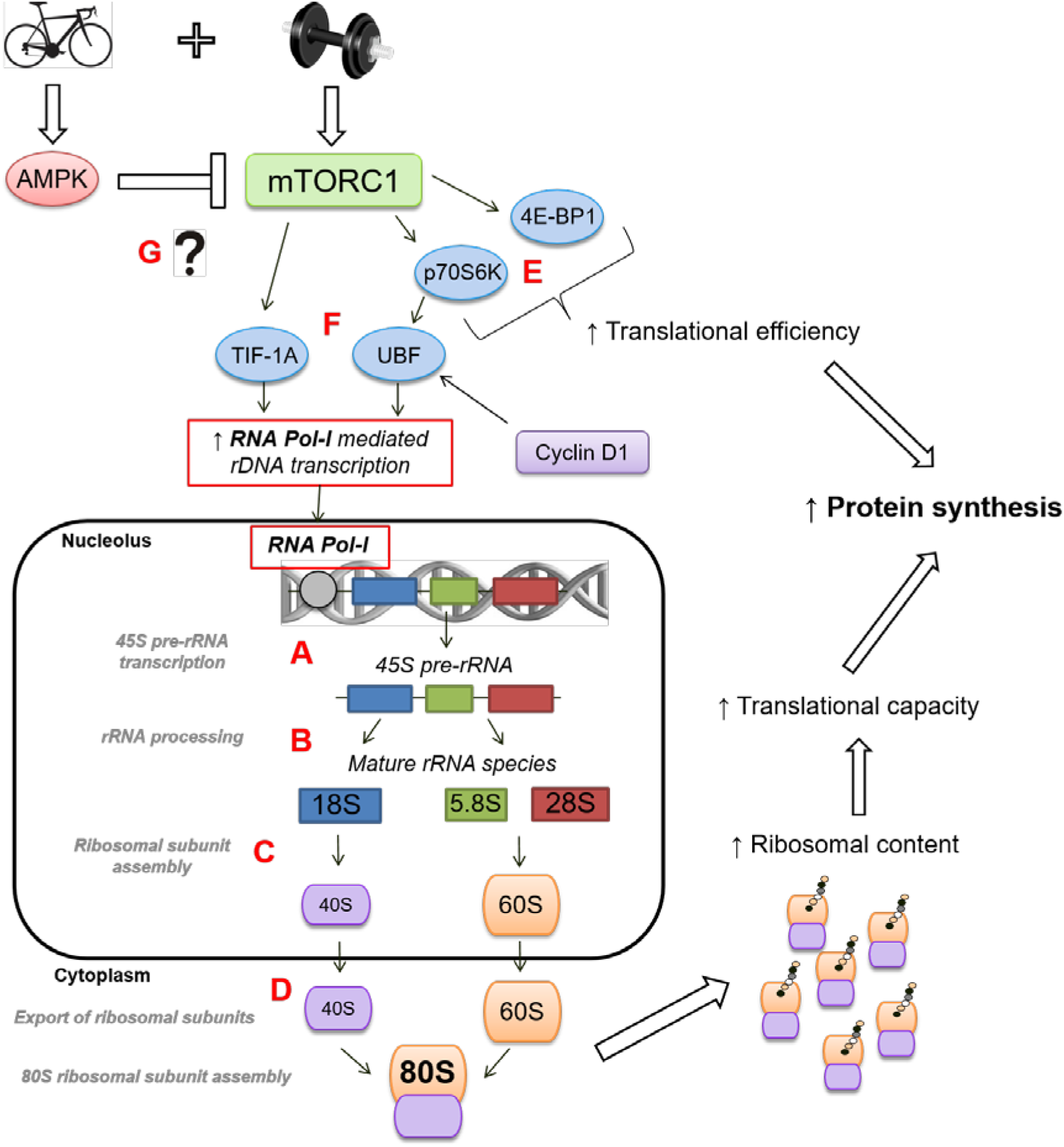
Overview of the role of mTORC1 signalling in promoting ribosome biogenesis following a single session of resistance exercise, and the potential effect of incorporating endurance training (i.e, performing concurrent training). Adapted from ^24^. Ribosome biogenesis involves transcription of the 45S rRNA (ribosomal RNA) precursor (45S pre-rRNA) **(A)** mediated by RNA Polymerase I (Pol-I), processing of the 45S pre-rRNA into several smaller rRNAs (18S, 5.8S and 28S rRNAs) **(B)**, assembly of these rRNAs and other ribosomal proteins into ribosomal subunits (40S and 60S) **(C)**, and nuclear export of these ribosomal subunits into the cytoplasm ^24,26^ **(D)**. As well as regulating translational efficiency via downstream control of p70S6K (p70 kDa ribosomal protein subunit kinase 1) and 4E-BP1 (eukaryotic initiation factor 4E binding protein 1) **(E)**, mTORC1 is a key mediator of ribosome biogenesis by regulating transcription factors for genes encoding RNA Pol-I (and also RNA Pol-II and nIII, which are not shown in figure) ^25^. Transcription of the 45S pre-rRNA by RNA Pol-I requires a transcriptional complex including TIF-1A (transcription initiation factor 1A; also known as RRN5) and UBF (upstream binding factor), both of which are regulated by the mTORC1 pathway ^28^,29 **(F)**. Activation of AMPK is known to inhibit mTORC1 signalling in rodent skeletal muscle ^64^, and AMPK activation in skeletal muscle is traditionally associated with endurance-type exercise. However, whether signalling events initiated by endurance training, when performed concurrently with resistance training, have the potential to interfere with mTORC1-mediated regulation of ribosome biogenesis is currently unclear **(G)**.

Studies in both human ^33-35^ and rodent skeletal muscle ^36-41^ suggest ribosome biogenesis, as indexed by increases in total RNA content (>85% of which comprises rRNA) ^24^, and increased mRNA expression of several RNA Pol-I regulatory factors, including UBF, cyclin D1 and TIF 1A, occurs concomitantly with muscle hypertrophy. In addition, attenuated rodent skeletal muscle hypertrophy with ageing ^35,42,43^ and rapamycin treatment ^40^ is associated with reduced markers of ribosome biogenesis, suggesting translational capacity is closely linked to the regulation of skeletal muscle mass. Despite the links between skeletal muscle hypertrophy and ribosome biogenesis ^24,33,34^, studies investigating molecular interference following concurrent exercise in human skeletal muscle have only measured transient (<6 h) post-exercise changes in translational efficiency (as indexed by mTORC1 signalling) and MPS ^9-18^. No studies have investigated changes in markers of ribosome biogenesis either after single bouts of concurrent exercise or following periods of concurrent training. Whether attenuated muscle hypertrophy following concurrent training could be explained, at least in part, by attenuated ribosome biogenesis is unknown.

The aim of this study was therefore to investigate changes in markers of ribosome biogenesis and mTORC1 signalling after eight weeks of concurrent training compared with resistance training undertaken alone. A secondary aim was to determine the potential role of endurance training intensity in modulating skeletal muscle ribosome biogenesis adaptation to concurrent training, by comparing concurrent training incorporating either high-intensity interval training (HIT) or moderate-intensity continuous training (MICT). The induction of these responses in skeletal muscle was also investigated following a single exercise session performed post training. It was hypothesised that compared with resistance training alone, concurrent training would attenuate the training-induced increase in markers of skeletal muscle ribosome biogenesis, and the induction of mTORC1 signalling, both at rest post-training and after a single training session performed in a training-accustomed state. It was further hypothesised that concurrent training incorporating HIT would preferentially attenuate training-induced skeletal muscle hypertrophy relative to resistance training alone, and this would be associated with an attenuation of markers of skeletal muscle ribosome biogenesis.

## 3. Results

### Training-induced changes in maximal strength and lean body mass

In brief, and as previously reported ^44^, one-repetition maximum (1-RM) leg press strength was improved for RT (mean change ±90% confidence interval; 38.5 ±8.5%; effect size [ES] ±90% confidence interval; 1.26 ±0.24; *P*<0.001), HIT+RT (28.7 ±5.3%; ES, 1.17 ±0.19; *P*<0.001) and MICT+RT (27.5 ±4.6%, ES, 0.81 ±0.12; *P*<0.001). Consistent with previous reports of interference to strength development with concurrent training ^3,19^, the magnitude of this change was greater for RT vs. both HIT+RT (7.4 ±8.7%; ES, 0.40 ±0.40) and MICT+RT (8.2 ±9.9%; ES, 0.60 ±0.45). Despite the differences in strength, lower-body lean mass was similarly increased for RT (4.1 ±2.0%; ES; 0.33 ±0.16; *P*=0.023) and MICT+RT (3.6 ±2.4%; ES; 0.45 ±0.30; *P*=0.052); however, this increase was attenuated for HIT+RT (1.8 ±1.6%; ES; 0.13 ±0.12; *P*=0.069).

### Physiological and psychological responses to the final training session

Heart rate and rating of perceived exertion (RPE)

During the final training session, there was a higher average heart rate (mean difference range, 14 ±12 to 19 ±14 bpm; ES, 1.04 ±0.88 to 1.22 ±0.89; *P* ≤ 0.043; Table 1) and rating of perceived exertion (RPE) (2 ±2 to 4 ±2 AU; ES, 1.51 ±0.86 to 2.15 ±0.87; *P* ≤ 0.06) for HIT compared with MICT. During the final training session, venous blood lactate (Table 1) was also higher for HIT compared with MICT at all time points both during cycling (mean difference range, 0.8 ±0.5 to 4.5 ±1.1 mmol·L^−1^; ES range, 1.46 ±0.87 to 3.65 ±0.85; *P* ≤ 0.01) and during the 15-min recovery period after cycling (3.5 ±1.0 to 5.0 ±1.2 mmol·L^−1^; ES, 3.11 ±0.85 to 3.68 ±0.85; *P* < 0.001). Venous blood glucose (Table 1) was also higher for HIT vs. MICT after 16, 22, 28 and 34 min cycling (0.4 ±0.7 to 1.6 ±0.9 mmol·L^−1^; ES, 0.54 ±0.86 to 1.52 ±0.86; *P* ≤ 0.039), and during the 15-min recovery period after cycling (0.9 ±0.7 to 1.8 ±1.0 mmol·L^−1^; ES, 1.11 ±0.85 to 1.50 ±0.85; *P* ≤ 0.041).

**Table 1.** Physiological and psychological (RPE) responses to a single bout of high-intensity interval training (HIT) or work-matched moderate-intensity continuous training (MICT) performed during the final training session.

|  | Time (min) |  |  |  |  |  |  |  |  |  |
| --- | --- | --- | --- | --- | --- | --- | --- | --- | --- | --- |
|  | Rest | 10 | 16 | 22 | 28 | 34 | +2 | +5 | +10 | +15 |
| Lactate (mmol·L<sup>-1</sup>) |  |  |  |  |  |  |  |  |  |  |
| HIT | 0.7 ± 0.3 | 2.6 ± 0.6 *# | 5.4 ± 1.4 *# | 6.8 ± 1.2 *# | 7.3 ± 1.4 *# | 7.3 ± 1.3 *# | 7.3 ± 1.8 *# | 7.2 ± 1.6 *# | 6.0 ± 1.5 *# | 4.9 ± 1.4 *# |
| MICT | 0.7 ± 0.3 | 1.7 ± 0.5 * | 2.6 ± 0.8 * | 2.7 ± 0.8 * | 2.8 ± 0.9 * | 2.8 ± 1.0 * | 2.4 ± 0.8 * | 2.2 ± 0.8 * | 1.8 ± 0.7 * | 1.4 ± 0.5 * |
| Glucose (mmol·L<sup>-1</sup>) |  |  |  |  |  |  |  |  |  |  |
| HIT | 4.7 ± 0.8 | 4.6 ± 0.9 | 4.8 ± 0.9 | 5.0 ± 0.9 # | 5.4 ± 1.1 # | 5.9 ± 1.2 *# | 6.3 ± 1.5 *# | 6.2 ± 1.3 *# | 5.9 ± 1.2 *# | 5.4 ± 1.0 # |
| MICT | 4.5 ± 0.5 | 4.5 ± 0.4 | 4.4 ± 0.6 | 4.2 ± 0.3 | 4.3 ± 0.4 | 4.3 ± 0.4 | 4.5 ± 0.5 | 4.7 ± 0.4 | 4.6 ± 0.4 | 4.5 ± 0.4 |
| Heart rate (beats·min<sup>-1</sup>) |  |  |  |  |  |  |  |  |  |  |
| HIT | 63 ± 11 | 154 ± 9 *# | 162 ± 9 *# | 166 ± 9 *# | 170 ± 10 *# | 173 ± 9 *# | - | - | - | - |
| MICT | 66 ± 5 | 140 ± 6 * | 147 ± 17 * | 150 ± 16 * | 152 ± 17 * | 154 ± 17 * | - | - | - | - |
| RPE (AU) |  |  |  |  |  |  |  |  |  |  |
| HIT | 6 ± 0 | 13 ± 3 * | 15 ± 3 *# | 17 ± 2 *# | 18 ± 2 *# | 18 ± 2 *# | - | - | - | - |
| MICT | 6 ± 0 | 11 ± 2 * | 12 ± 2 * | 13 ± 2 * | 14 ± 2 * | 14 ± 2 * | - | - | - | - |

Values are means ± SD. HIT, high-intensity interval training cycling; MICT, continuous cycling; RPE, rating of perceived exertion. \*, $P < 0.05$ vs. rest; #, $P < 0.05$ vs. MICT at same time point.

After completion of RE in the final training session, venous blood lactate (Table 2) was higher for HIT+RT vs. RT after 0, 2, 5, 10, 60, 90 and 180 min of recovery (0.1 ±0.1 to 1.4 ±0.9 mmol·L^−1^; ES, 0.80 ±0.84 to 1.74 ±0.84; *P* ≤ 0.095), and higher for HIT+RT vs. MICT+RT at all timepoints (0.1 ±0.1 to 1.1 ±1.4 mmol·L^−1^; ES, 0.73 ±0.87 to 1.82 ±0.86; *P* ≤ 0.161). Post-RE venous blood glucose (Table 2) was lower for HIT+RT vs. RT after 2, 10, and 30 min of recovery (0.3 ±0.2 to 0.3 ±0.3 mmol·L^−1^; ES, −0.65 ±0.84 to −1.02 ±0.84; *P* ≤ 0.193), and higher for HIT+RT vs. RT after 60 min of recovery (0.4 ±0.4 mmol·L^−1^; ES, 0.88 ±0.84; *P* = 0.077). Blood glucose was higher for MICT vs. HIT+RT at +30 min of recovery (0.3 ±0.2 mmol·L^−1^; ES, 1.29 ±0.86; *P* = 0.021), and lower for HIT+RT vs. MICT+RT at +60 min of recovery (0.2 ±0.2 mmol·L^−1^; ES, −1.09 ±0.85; *P* = 0.045).

**Table 2.** Venous blood lactate and glucose responses to a single bout of resistance exercise (RE) either performed alone (RT) or when performed after either high-intensity interval training (HIT+RT) or work-matched moderate-intensity continuous training (MICT+RT) during the final training session.

|  | Time (min) |  |  |  |  |  |  |  |
| --- | --- | --- | --- | --- | --- | --- | --- | --- |
|  | End | +2 | +5 | +10 | +30 | +60 | +90 | +180 |
| Lactate (mmol·L<sup>-1</sup>) |  |  |  |  |  |  |  |  |
| RT | 2.1 ± 0.7 * | 2.3 ± 0.9 * | 2.2 ± 1.0 * | 1.7 ± 0.8 * | 1.3 ± 1.3 | 0.7 ± 0.3 | 0.6 ± 0.2 | 0.5 ± 0.2 |
| HIT+RT | 3.5 ± 1.3 *‡ | 3.6 ± 1.5 * | 3.3 ± 1.4 * | 2.6 ± 1.2 * | 1.6 ± 0.4 *# | 1.2 ± 0.3 *#‡ | 0.8 ± 0.1 #‡ | 0.7 ± 0.1 |
| MICT+RT | 2.4 ± 1.2 * | 2.5 ± 1.4 * | 2.2 ± 1.2 * | 1.7 ± 0.7 * | 0.9 ± 1.3 | 0.7 ± 0.2 | 0.6 ± 0.1 | 0.5 ± 0.2 |
| Glucose (mmol·L<sup>-1</sup>) |  |  |  |  |  |  |  |  |
| RT | 4.7 ± 0.3 | 4.7 ± 0.4 | 4.7 ± 0.4 | 4.7 ± 0.4 | 4.7 ± 0.3 ^ | 4.3 ± 0.5 | 4.5 ± 0.3 | 4.5 ± 0.2 |
| HIT+RT | 4.5 ± 0.9 | 4.5 ± 0.4 | 4.5 ± 0.4 | 4.4 ± 0.4 | 4.5 ± 0.2 | 4.7 ± 0.3 # | 4.5 ± 0.2 | 4.6 ± 0.3 |
| MICT+RT | 4.6 ± 0.3 | 4.6 ± 0.3 | 4.7 ± 0.2 | 4.6 ± 0.2 | 4.7 ± 0.2 ^ | 4.4 ± 0.1 | 4.4 ± 0.2 | 4.4 ± 0.4 |

Values are means ± SD. HIT+RT, high-intensity interval training cycling and resistance training; MICT+RT, continuous cycling and resistance training; RT, resistance training; \*, $P < 0.05$ vs. rest; #, $P < 0.05$ vs. MICT at same time point; ^, $P < 0.05$ vs. HIT at same time point.; ‡, $P < 0.05$ vs. RT at same time point.

### Training-induced changes in markers of ribosome biogenesis

The total RNA content of skeletal muscle was measured as an index of translational capacity (and hence ribosomal content), since ribosomal RNA comprises over 85% of the total RNA pool ^45^. Pre-training total RNA content was higher for the RT group vs. both HIT+RT (38 ±17%; ES, −1.48 ±0.84; *P* = 0.005; Table 3) and MICT+RT (25 ±12%; ES, 1.47 ±0.85; *P* = 0.010). Total RNA content was decreased in the basal state post-training in the RT group (−11 ±5%; ES, −0.17 ±0.09; *P* = 0.025), and was not substantially changed for either HIT+RT (32 ±18%; ES, 0.30 ±0.15; *P* = 0.077) or MICT+RT (20 ±15%; ES, 0.12 ±0.08; *P* = 0.083). The change in total RNA content between pre-and post-training was, however, greater for both HIT+RT (48 ±39%; ES, 1.14 ±0.76) and MICT+RT (34 ±24%; ES, 1.24 ±0.75) vs. RT.

**Table 3.** Total RNA content and type I and type II muscle fibre cross-sectional area (CSA) of the vastus lateralis before (PRE-T) and after (POST-T) eight weeks of either RT alone, or RT combined with either high-intensity interval training (HIT+RT) or moderate-intensity continuous training (MICT+RT).

| Measure | PRE-T | POST-T |
| --- | --- | --- |
| Total skeletal muscle RNA (ng/mg tissue) |  |  |
| RT | 914 ± 202<sup>^</sup> | 810 ± 134<sup>*</sup> |
| HIT+RT | 581 ± 176 | 740 ± 129 |
| MICT+RT | 680 ± 81 | 818 ± 133 |
| Type I muscle fibre CSA (µm<sup>2</sup>) |  |  |
| RT | 4539 ± 848 | 5533 ± 1913<sup>*b</sup> |
| HIT+RT | 6713 ± 1849 | 5183 ± 1413 |
| MICT+RT | 5509 ± 2326 | 5228 ± 1277 |
| Type II muscle fibre CSA (µm<sup>2</sup>) |  |  |
| RT | 5296 ± 1347 | 6456 ± 2235 |
| HIT+RT | 6470 ± 1481 | 6621 ± 2018 |
| MICT+RT | 5051 ± 1531 | 5728 ± 688 |
| Data presented are means ± SD. * = $P < 0.05$ vs. PRE-T, ^ = $P < 0.05$ vs. both HIT+RT and MICT+RT at PRE-T, b = change between PRE-T and POST-T substantially greater vs. HIT+RT. | | |

Given the observed changes in skeletal muscle RNA content, we also investigated training induced changes in components of the ribosomal machinery in skeletal muscle, including expression levels of the 45S rRNA precursor, and the mature forms of the 5.8S, 18S and 28S rRNAs.

Expression of 45S pre-rRNA was unaltered by training for all groups (Figure 2); however, greater training-induced changes in 45S pre-rRNA expression were noted for both HIT+RT (58 ±76%; ES, 0.71 ±0.71) and MICT+RT (75 ±81%; ES, 0.85 ±0.68) vs. RT. There were no substantial changes, nor between-group differences, in 45S pre-rRNA expression during the final training session for either training group.

**Figure 2.**
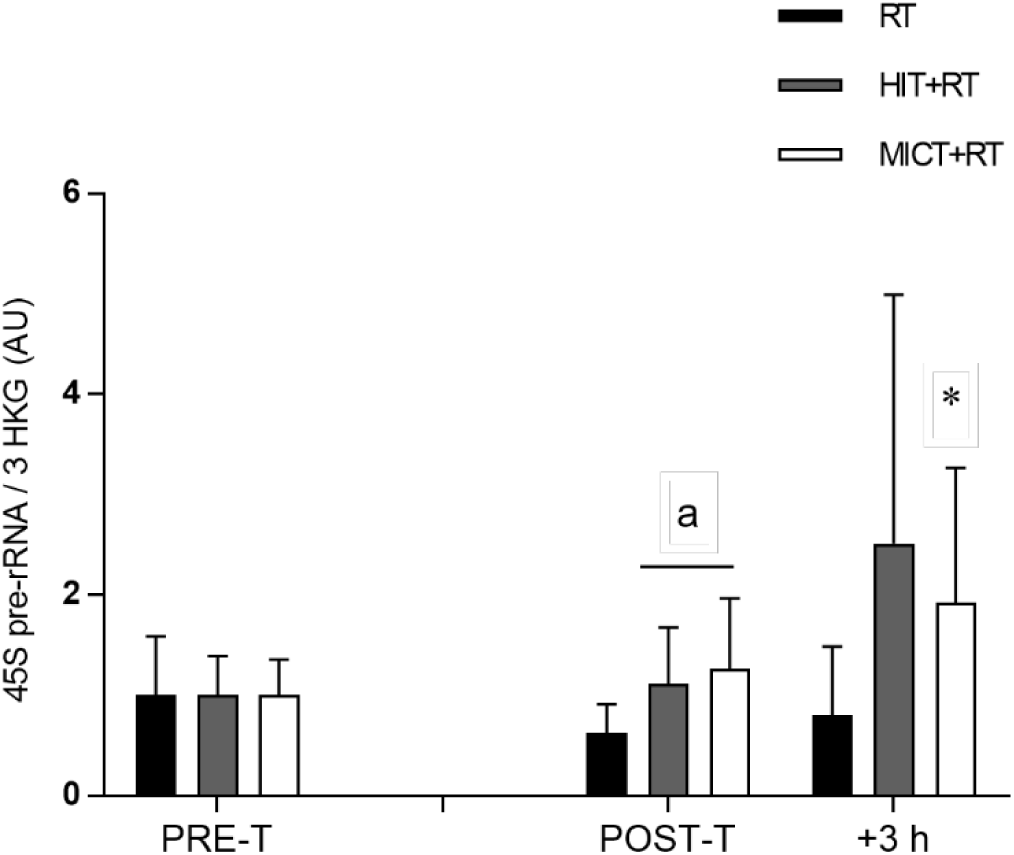
Expression of 45S pre-rRNA relative to the geometric mean of the expression of three housekeeping genes (HKG) (cyclophillin, β2M and TBP) before (PRE-T) and after (POST-T) eight weeks of either RT alone, or RT combined with either high-intensity interval training (HIT+RT) or moderate-intensity continuous training (MICT+RT), and 1 h and 3 h after a single exercise bout performed post-training. Data presented are means ± SD and expressed relative to the PRE-T value for each corresponding group. * = *P* < 0.05 vs. PRE-T, a = change between PRE-T and POST-T substantially different vs. RT.

We used specifically-designed primers ^34^ to distinguish between the expression levels of mature rRNA species [designated as 5.8S, 18S and 28S (mature) rRNAs] and those transcripts still bound to the 45S rRNA precursor and hence indicative only of changes in 45S pre-rRNA expression [designated as 5.8S, 18S and 28S (span) rRNAs].

Expression of 5.85S rRNA (mature) was lower post-training for RT (−51 ±16%; ES, −0.69 ±0.31; *P* = 0.017; Figure 3A). Both HIT+RT (125 ±109%; ES, 1.27 ±0.73) and MICT+RT (120 ±111%; ES, 0.99 ±0.61) induced greater post-training increases in 5.8S rRNA (mature) expression vs. RT. Neither training group induced substantial post-exercise changes in 5.8S rRNA (mature) expression during the final training session. Expression of 5.8S rRNA (span) was also lower post-training for RT (−36 ±15%; ES, −0.51 ±0.27; *P* = 0.027; Figure 3B), and the basal training-induced change in 5.8S rRNA (span) expression was greater for HIT+RT vs. RT (112 ±116%; ES, 1.40 ±0.97).

**Figure 3.**
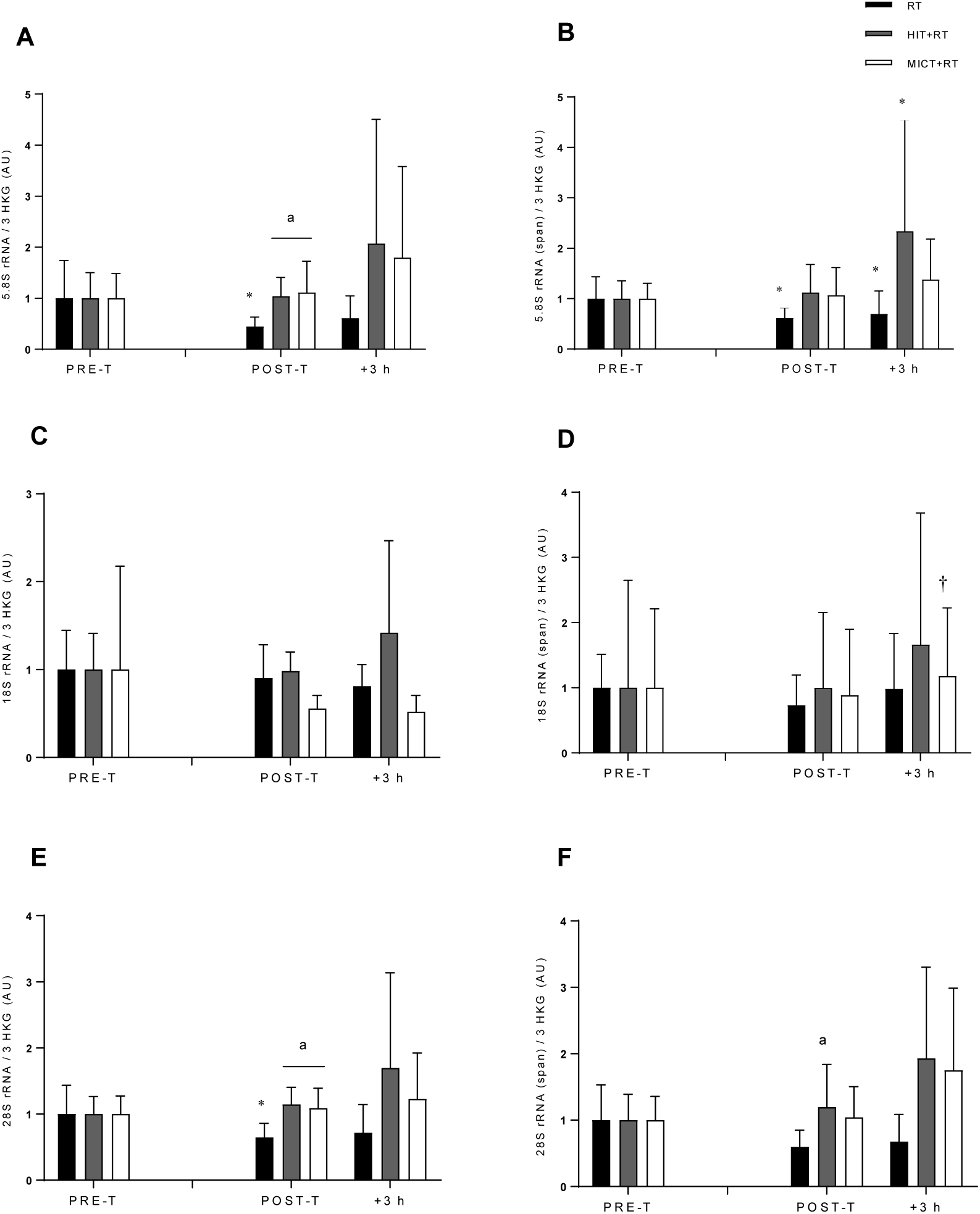
Expression of the mature rRNA transcripts 5.8S rRNA (A), 18S rRNA (C), and 28S rRNA (E), and rRNA transcripts bound to the 45S pre-RNA precursor: 5.8S rRNA (span) (B) 18S rRNA (span) (D) and 28S rRNA (span) (F) relative to the geometric mean of the expression of three housekeeping genes (HKG) (cyclophillin, β2M and TBP) before (PRE-T) and after (POST-T) eight weeks of either RT alone, or RT combined with either high-intensity interval training (HIT+RT) or moderate-intensity continuous training (MICT+RT), and 1 h and 3 h after a single exercise bout performed post-training. Data presented are means ± SD and expressed relative to the PRE-T value for each corresponding group. * = *P* < 0.05 vs. PRE-T, † = *P* < 0.05 vs. POST-T, a = change between PRE-T and POST-T substantially greater vs RT.

Expression of 18S rRNA (mature) was not substantially different at any time point, nor were there any substantial between-group differences in changes in 18S rRNA (mature) expression (Figure 3C). There were also no substantial effects of training or any between-group differences in changes in 18S rRNA (span) expression (Figure 3D), although increased 18S rRNA (span) expression was noted at +3 h during the final training bout for MICT+RT (63 ±48%; ES, 0.21 ±0.12; *P* = 0.029).

Resting levels of 28S rRNA (mature) expression were reduced post-training for RT (−33 ±15%; ES, −0.49 ±0.28; *P* = 0.037; Figure 3E). Greater training induced changes in basal 28S rRNA expression were noted for both HIT+RT (73 ±56%; ES, 1.23 ±0.71; *P* = 0.007) and MICT+RT (63 ±55%; ES, 1.10 ±0.74; *P* = 0.023) vs. RT. Neither training group induced substantial post exercise changes in 28S rRNA expression during the final training session. No changes in resting 28S rRNA (span) expression were noted in response to training for either group (Figure 3F). However, HIT+RT induced greater training-induced changes in basal 28S rRNA (span) expression compared with RT (123 ±109%; ES, 0.81 ±0.48).

### Ribosome biogenesis-related signalling responses

To determine potential upstream molecular events associated with changes in markers of ribosome biogenesis with concurrent versus single-mode resistance training, we investigated the regulation of key proteins (TIF-1A, UBF and cyclin D1) involved in promoting 45S rRNA precursor expression (Figure 1).

Post-training, basal levels of TIF-1A^Ser649^ phosphorylation were increased only for HIT+RT (133 ±102%; ES, 0.62 ±0.31; *P* = 0.047; Figure 4A). During the final training session, only RT was sufficient to increase TIF-1A phosphorylation at both +1 h (123 ±79%; ES, 0.45 ±0.19; *P* = 0.002) and +3 h (241 ±315%; ES, 0.69 ±0.46; *P* = 0.017), and this increase (between POST T to +3 h) was greater than for both HIT+RT (52 ±46%; ES, 0.76 ±0.89) and MICT+RT (75 ±24%; ES, 1.31 ±0.80).

**Figure 4.**
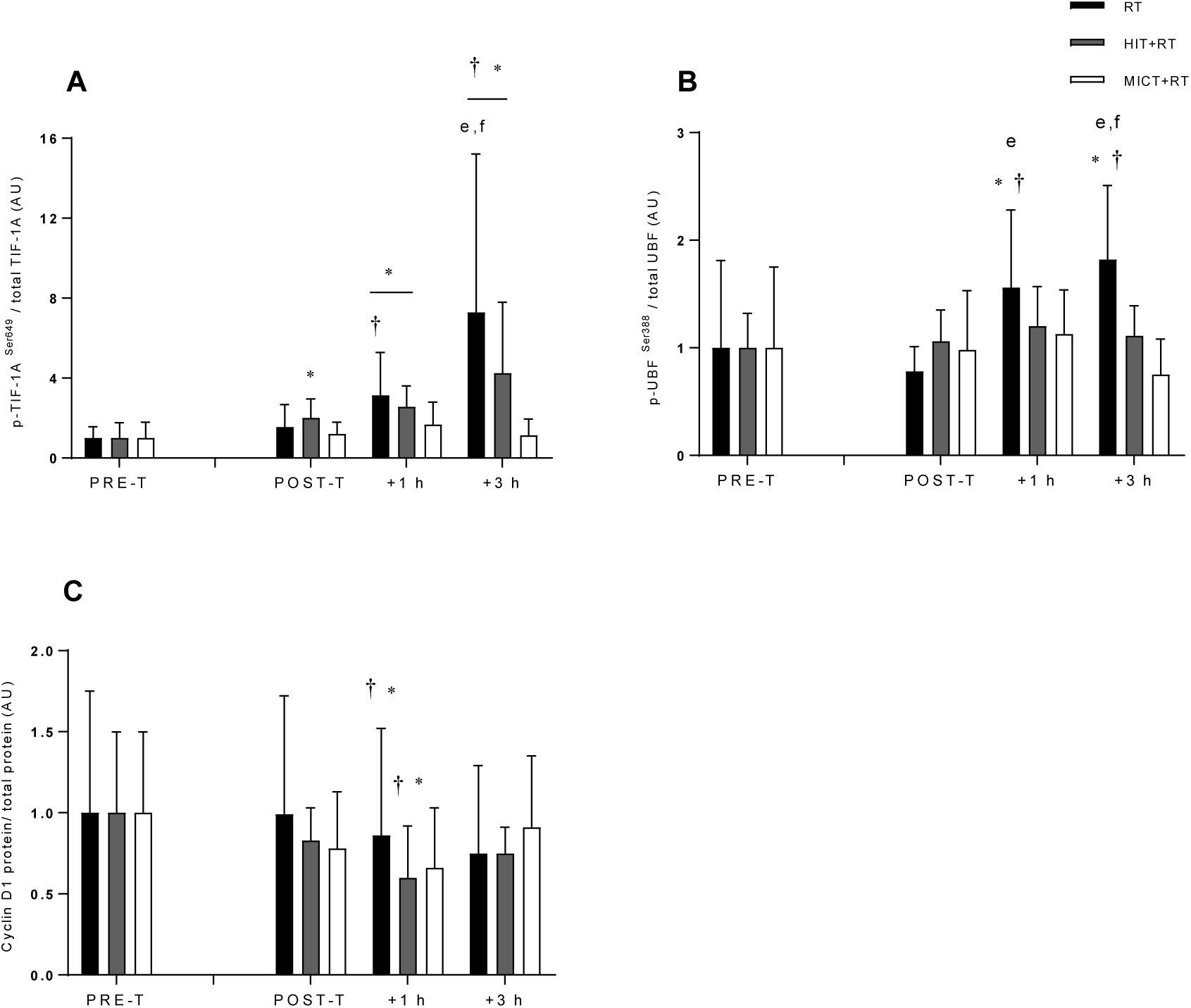
Phosphorylation of TIF-1A^Ser649^ (A), UBF^Ser388^ (B), and total protein content of cyclin D1 (C) before (PRE-T) and after (POST-T) eight weeks of either RT alone, or RT combined with either high-intensity interval training (HIT+RT) or moderate-intensity continuous training (MICT+RT), and 1 h and 3 h after a single exercise bout performed post-training. Data presented are means ± SD and expressed relative to the PRE-T value for each corresponding group. * = *P* < 0.05 vs. PRE-T, † = *P* < 0.05 vs. POST-T. Change from POST-T substantially greater vs. e = HIT+RT, f = MICT+RT.

A similar pattern was observed for UBF^Ser388^ phosphorylation, although no training group showed altered UBF^Ser388^ phosphorylation in the basal state post-training (see Figure 4B). As observed with TIF-1A, only RT was sufficient to increase UBF phosphorylation during the final training session at both +1 h (78 ±58%; ES, 0.82 ±0.45; *P* = 0.010) and + 3 h (125 ±72%; ES, 1.15 ±0.45; *P* = 0.001). RT also induced greater changes in UBF phosphorylation during the final training session at +1 h vs. both HIT+RT (32 ±23%; ES, 0.54 ±0.46) and MICT+RT (37 ±27%; ES, 0.61 ±0.55), and also at +3 h vs.both HIT+RT (49 ±17%; ES, 0.92 ±0.45) and MICT+RT (64 ±12%; ES, 1.35 ±0.42).

As observed for UBF, the protein content of cyclin D1 was unchanged between pre- and post training for all training groups (Figure 4C). However, a post-exercise reduction in cyclin D1 protein content was observed for HIT+RT at +1 h during the final training session (−34 ±7%; ES, −0.66 ±0.16; *P* = 0.008).

### mRNA responses related to ribosome biogenesis

We also measured the mRNA levels of select genes (TIF-1A, UBF, POLR1B, and cyclin D1) involved in promoting 45S rRNA precursor expression (Figure 1).

Neither training group altered basal TIF-1A mRNA content post-training (Figure 5A). During the final training session, only RT (26 ±12%; ES, 0.53 ±0.21; *P* = 0.003) and MICT+RT (36 ±35%; ES, 0.59 ±0.50; *P* = 0.038) increased TIF-1A mRNA expression at +3 h.

**Figure 5.**
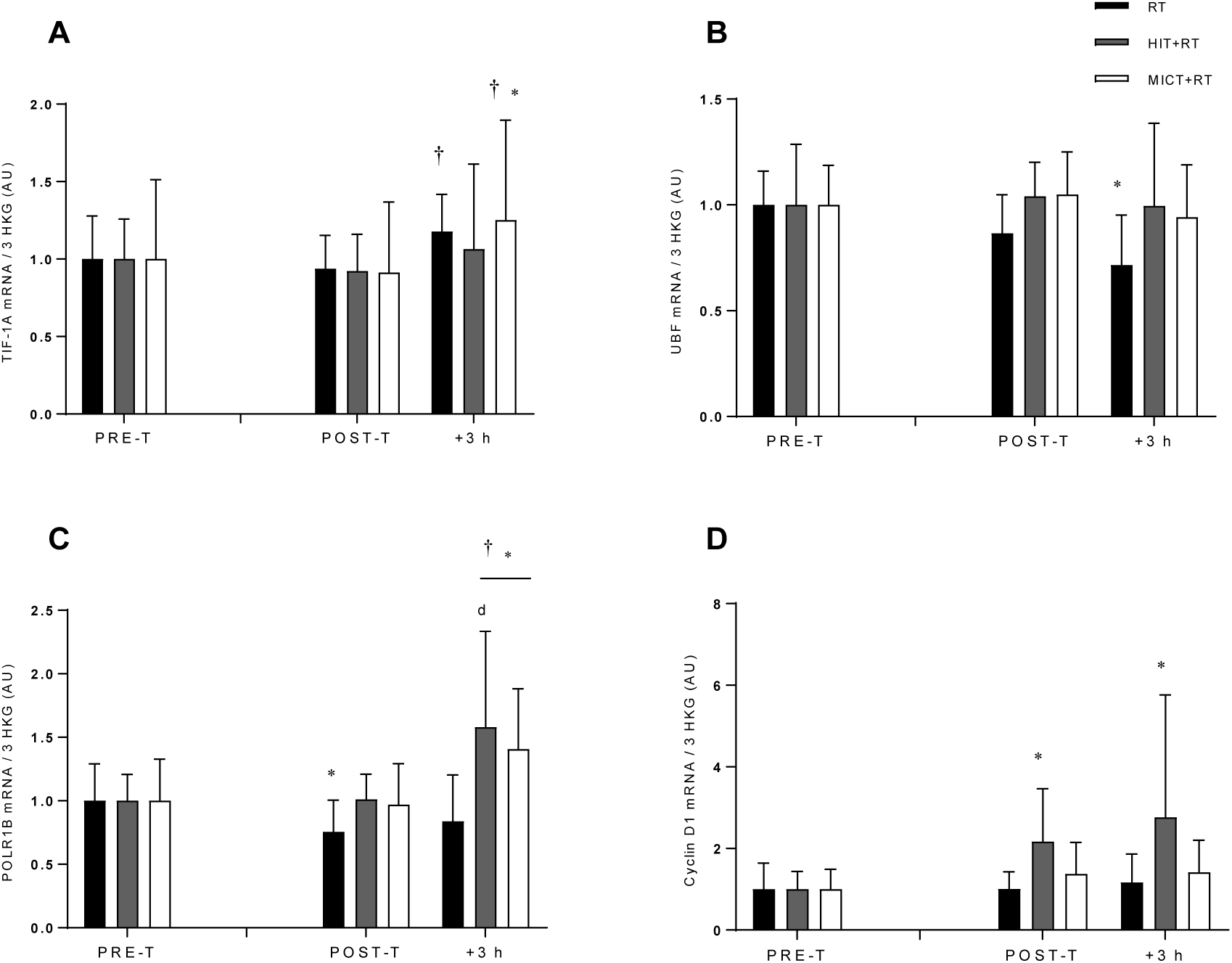
mRNA expression of TIF-1A (A), UBF (B), POLR1B (C), and cyclin D1 (D) relative to the geometric mean of the expression of three housekeeping genes (HKG) (cyclophillin, β2M and TBP) before (PRE-T) and after (POST-T) eight weeks of either RT alone, or RT combined with either high-intensity interval training (HIT+RT) or moderate-intensity continuous training (MICT+RT), and 1 h and 3 h after a single exercise bout performed post-training. Data presented are means ± SD and expressed relative to the PRE value for each corresponding group. * = *P* < 0.05 vs. PRE-T, † = *P* < 0.05 vs. POST-T. Change from POST-T substantially greater vs. d = RT.

Basal levels of UBF mRNA post-training were not altered by either training group (Figure 5B). There were no substantial changes in UBF expression during the final training session for either training group.

Basal expression of POLR1B mRNA was reduced post-training by RT alone (−26 ±16%; ES, - 0.44 ±0.32; *P* = 0.026; Figure 5C). Only HIT+RT (44 ±42%; ES, 0.57 ±0.44; *P* = 0.047) and MICT+RT (48 ±43%; ES, 0.51 ±0.37; *P* = 0.033) increased POLR1B mRNA expression at + 3 during the final training session.

Post-training basal expression of cyclin D1 mRNA was increased only for HIT+RT (101 ±54%; ES, 0.59 ±0.22; *P* = 0.001; Figure 5D). No post-exercise changes in cyclin D1 mRNA were noted in response to the final training session for either training group.

### AMPK/mTORC1-related signalling responses

Given the observed changes in TIF-1A and UBF phosphorylation following training, we investigated changes in mTORC1 signalling as a potential upstream regulator of these responses ^25^, as well as AMPK signalling as a negative regulator of mTORC1 signalling ^7^.

The phosphorylation of AMPK^Thr172^ was unchanged in the basal state post-training for all groups (Figure 6A). AMPK phosphorylation was, however, increased by RT at +1 h during the final training session (78 ±72%; ES, 0.34 ±0.23; *P* = 0.031). RT also induced a greater change in AMPK phosphorylation at +3 h during the final training session vs. MICT+RT (59 ±44%; ES, 0.79 ±0.83), but not vs. HIT+RT (54 ±49%; ES, 0.69 ±0.83).

**Figure 6.**
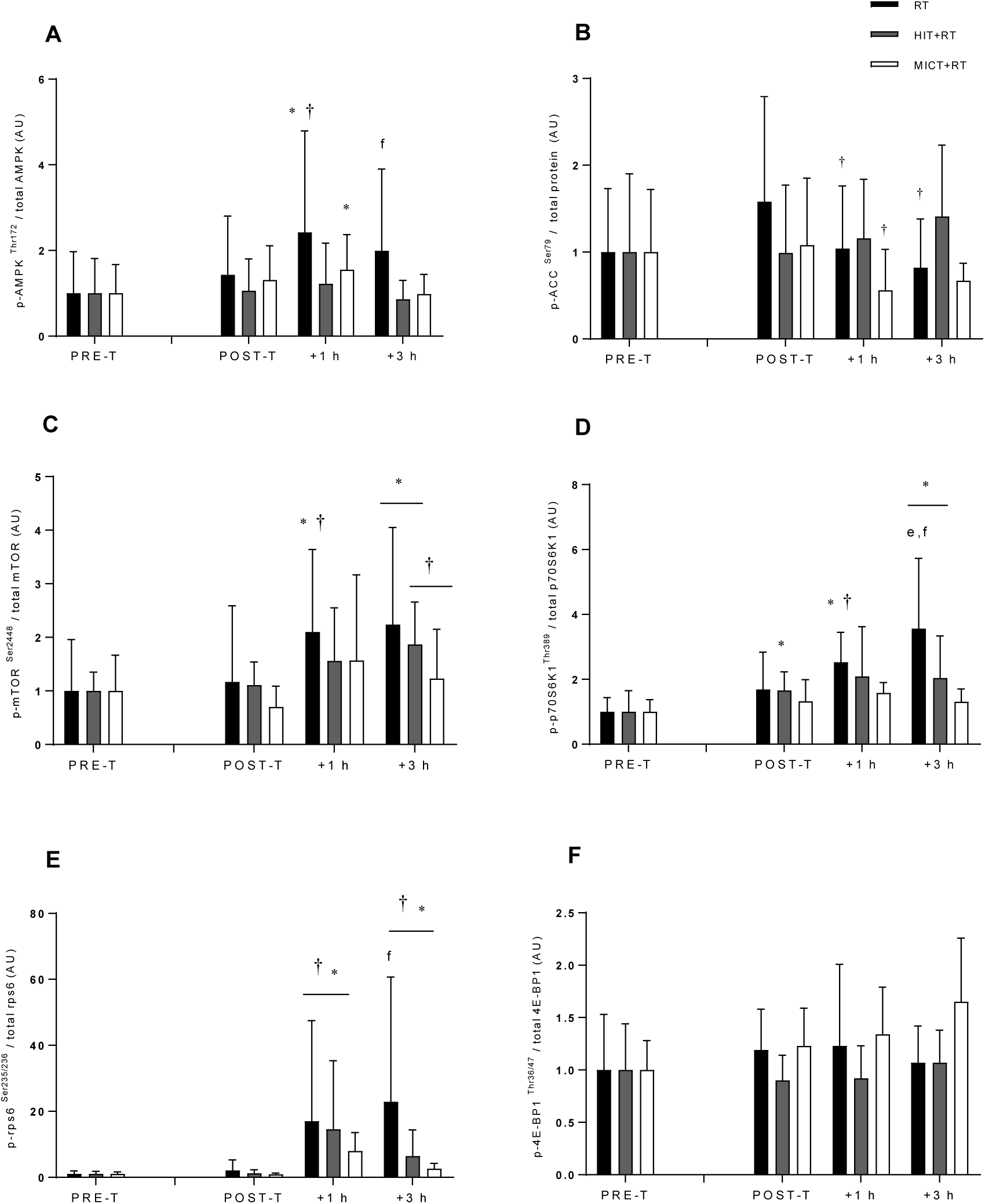
Phosphorylation of AMPK^Thr172^ (A), ACC^Ser79^ (B), mTOR^Ser2448^ (C), p70S6K^Thr389^ (D), rps6^Ser235/236^ (E) and 4E-BP1^Thr36/47^ (F) before (PRE-T) and after (POST-T) eight weeks of either RT alone, or RT combined with either high-intensity interval training (HIT+RT) or moderate-intensity continuous training (MICT+RT), and 1 h and 3 h after a single exercise bout performed post-training. Data presented are means ± SD and expressed relative to the PRE value for each corresponding group. * = *P* < 0.05 vs. PRE-T, † = *P* < 0.05 vs. POST-T. Change from POST-T substantially greater vs. e = HIT+RT, f = MICT+RT.

As observed with AMPK^Thr172^ phosphorylation, neither training group had altered ACC^Ser79^ phosphorylation in the basal state post-training (Figure 6B). However, reductions in ACC phosphorylation were noted at +1 h in the final training session for both RT (−36 ±22%; ES, - 0.28 ±0.20; *P* = 0.026) and MICT+RT (−46 ±20%; ES, −0.56 ±0.33; *P* = 0.016), and at +3 h for RT (−45 ±20%; ES, −0.37 ±0.22; *P* = 0.012). Compared with RT, HIT+RT induced greater changes in ACC phosphorylation during the final training session at both +1 h (99 ±100%; ES, 0.65 ±0.46) and +3 h (169 ±168%; ES, 0.94 ±0.56).

As observed with both AMPK^Thr172^ and ACC^Ser79^, the phosphorylation of mTOR^Ser2448^ was unchanged in the basal state post-training for all training groups (Figure 6C). In response to the final training bout, mTOR phosphorylation was increased at +1 h only for RT (105 ±137%; ES, 0.46 ±0.40; *P* = 0.048), and not for either HIT+RT (30 ±71%; ES, 0.32 ±0.62; *P* = 0.320) or MICT+RT (77 ±184%; ES, 0.37 ±0.59; *P* = 0.218), and was increased at +3 h only for HIT+RT (70 ±45%; ES, 0.64 ±0.31; *P* = 0.030).

The observed changes in p70S6K1^Thr389^ phosphorylation mirrored changes in mTOR^Ser2448^ phosphorylation (Figure 6D). Indeed, only RT was sufficient to increase p70S6K1^Thr389^ phosphorylation during the final training session at +1 h (78 ±77%; ES, 0.51 ±0.37; *P* = 0.026). RT alone also induced a greater change in p70S6K1 phosphorylation at +3 h compared with both HIT+RT (47 ±50%; ES, 0.86 ±1.13) and MICT+RT (50 ±46%; ES, 0.88 ±1.05).

All training groups increased rps6^Ser235/236^ phosphorylation during the final training bout at +1 h (RT: 700 ±678%; ES, 0.75 ±0.28; *P* < 0.001; HIT+RT: 475 ±572%; ES, 0.66 ±0.33; *P* = 0.005; MICT+RT: 621 ±420%; ES, 1.49 ±0.42; *P* < 0.001) and +3 h (RT: 967 ±1047%; ES, 0.85 ±0.31; *P* < 0.001; HIT+RT: 294 ±319%; ES, 0.51 ±0.28; *P* = 0.006; MICT+RT: 176 ±200%; ES, 0.76 ±0.51; *P* = 0.026; Figure 6E). The change in rps6 phosphorylation at +3 h was, however, greater for RT vs. MICT+RT (74 ±29%; ES, 0.72 ±0.51) but not vs. HIT+RT (63 ±41%; ES, 0.57 ±0.56).

Despite the evidence of increased mTORC1 signalling as indexed via enhanced p70S6K1^Thr389^ phosphorylation, we observed no between-group differences in 4E-BP1^Thr36/47^ phosphorylation at any time point (Figure 6F).

### Muscle fibre CSA responses

We also performed immunohistochemical analyses to determine fibre-type specific changes in muscle fibre CSA induced by the concurrent versus single-mode resistance training protocols.

Type I muscle fibre CSA (see Table 3) was increased by RT alone (15 ±13%; ES, 0.10 ±0.08; *P* = 0.035), but not for either HIT+RT (−23 ±19%; ES, −0.09 ±0.08; *P* = 0.135) or MICT+RT (0.4 ±17%; ES, 0.00 ±-0.14; *P* = 0.989). The training-induced change in type I fibre CSA was also greater for RT compared with HIT+RT (34 ±22%; ES, 1.03 ±0.80), but not MICT+RT (15 ±54%; ES, 0.39 ±1.45).

Type II muscle fibre CSA (see Table 3) was not substantially altered by either RT alone (19 ±27%; ES, 0.09 ±0.12; *P* = 0.139), HIT+RT (0.4 ±24%; ES, 0.00 ±0.08; *P* = 0.974) or MICT+RT (16 ±14%; ES, 0.19 ±0.16; *P* = 0.344). Representative immunohistochemical images are shown in Figure 7.

**Figure 7.**
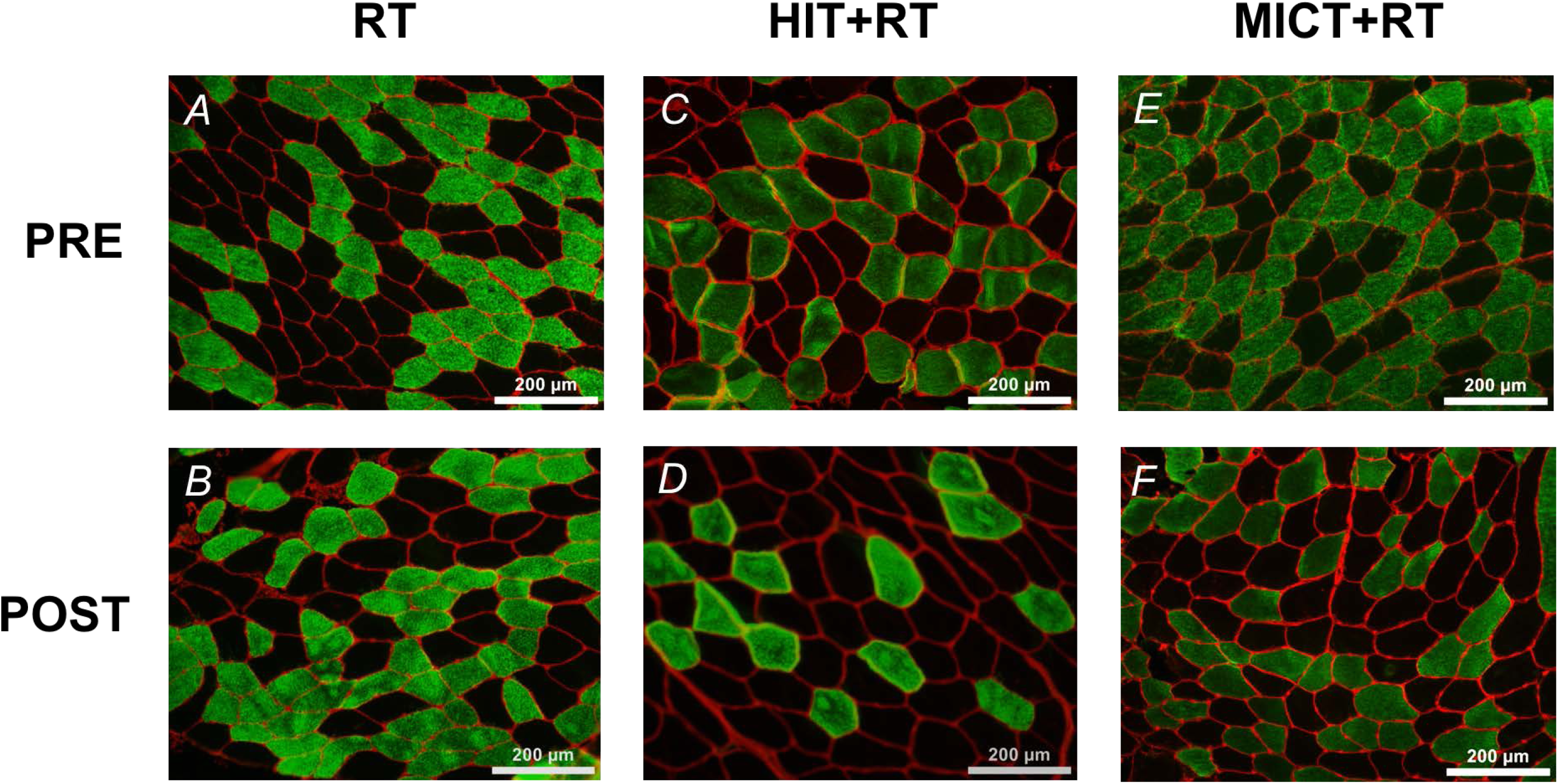
Representative immunohistochemical images of muscle cross-sections obtained before (PRE) and after (POST) eight weeks of either RT alone (images A and B, respectively), or RT combined with either high-intensity interval training (HIT+RT; images C and D, respectively) or moderate-intensity continuous training (MICT+RT; images E and F, respectively). Muscle fibre membranes are stained red, type I muscle fibres are stained green, and type II muscle fibres are unstained.

## 4. Discussion

Previous investigations on molecular responses and adaptations in skeletal muscle to concurrent training have focused almost exclusively on markers of post-exercise translational efficiency (i.e., mTORC1 signalling and rates of MPS)^9-18^. For the first time, we present data on the regulation of translational capacity (i.e., ribosome biogenesis) in skeletal muscle with concurrent training compared with resistance training performed alone. The major findings were that training-induced changes in markers of ribosome biogenesis, including total RNA content and expression of some mature rRNA species (i.e., 5.8S and 28S, but not 18S) were more favourable following concurrent training compared with resistance training alone, and irrespective of the endurance training intensity employed. These responses occurred despite a single bout of resistance exercise, when performed post-training, further inducing both mTORC1- and ribosome biogenesis-related signalling (i.e., TIF-1 and UBF phosphorylation) compared with concurrent exercise. These observations also contrasted with our findings regarding changes in muscle fibre-type specific hypertrophy, which was greater in type I muscle fibres for the resistance training group, suggesting a disconnect between training induced changes in markers of ribosome biogenesis and muscle fibre hypertrophy.

To investigate the effects of concurrent versus single-mode resistance training on markers of skeletal muscle ribosome biogenesis, we measured training-induced changes in total RNA content and basal expression of mature ribosome species 5.8S, 18S, and 28S, as well as early post-exercise changes in mature rRNA expression. Contrary to our hypothesis, resistance training alone induced small decreases in the levels of both the 5.8S and 28S rRNAs in the basal state post-training, while the training-induced change in both of these mature rRNA species was greater with concurrent exercise compared with resistance training alone. Neither training protocol induced any changes in 18S rRNA expression. Previous work in humans has observed basal increases in 5.8S, 18S, and 28S rRNA expression in human skeletal muscle after 8 weeks of resistance training, all of which were reduced 1 h following a single session of resistance exercise performed post-training ^34^. The present data contrast with these findings by suggesting that resistance training performed alone was an insufficient stimulus to increase mature rRNA content, whereas concurrent exercise was sufficient to increase mature 5.8S and 28S expression after a single post-training exercise bout.

Consistent with the training-induced changes in both 5.8S and 28S rRNA expression with resistance training performed alone, a small reduction in basal total RNA content in skeletal muscle was observed within this cohort. Despite this paradoxical finding, it is interesting to note total RNA content was higher at PRE-T for the RT group compared with both the HIT+RT and MICT groups (1.6- and 1.3-fold, respectively). The reason for this between-group discrepancy at baseline is not immediately clear, given we previously showed no differences in baseline lean mass measured via DXA or lower-body 1-RM strength in these participants ^44^, suggesting other factors may have influenced the between-group differences in baseline skeletal muscle RNA content. It is also possible that the training program provided an insufficient stimulus to at least maintain this elevated basal RNA content for the RT group. Studies demonstrating robust increases in total RNA content concomitantly with rodent skeletal muscle hypertrophy typically employ supraphysiological methods for inducing muscle hypertrophy, such as synergist ablation ^36^,39,46,47, a stimulus that is clearly not replicated by resistance training in human models. Participant training status may also impact upon training induced changes in ribosome biogenesis in humans. The participants in the present study were actively engaging in resistance and/or endurance training for at least 1 year prior to commencing the study, suggesting a higher training status compared with those of Figueiredo et al. ^34^ who were likely untrained (although this was not made explicitly clear) and asked to refrain from resistance training for 3 weeks prior to the study ^34^. It is also possible that between group differences in training volume, which was clearly higher for the concurrent training groups compared with the RT group, may have impacted upon the training-induced changes in total skeletal muscle RNA content.

Despite the observed changes in skeletal muscle RNA content, resistance training alone was sufficient to increase type I, but not type II, muscle fibre CSA. The lack of any substantial type II fibre hypertrophy is likely due, at least in part, to the specific nature of the resistance training program employed, which was perhaps better-oriented for enhancing maximal strength rather than lean mass ^44^. Indeed, previously-published data indicates that the resistance training protocol employed in the present study was effective in improving maximal strength and measures of lean mass ^44^, although these changes did not transfer to detectable type II fibre hypertrophy. Nevertheless, in agreement with previous research ^2^,4, the training-induced increase in type I muscle fibre CSA was attenuated with concurrent exercise, albeit only when incorporating HIT, compared with resistance training performed alone. Despite these between group differences in fibre-type specific hypertrophy, we could find no evidence that the training-induced changes in lean mass or muscle fibre CSA were correlated with changes in total RNA content of skeletal muscle (data not shown). The apparent disconnect between training-induced changes in total RNA content and markers of muscle hypertrophy, both at the whole-body and muscle-fibre levels, suggests further investigation is required into relationship between changes in translational capacity and resistance training-induced hypertrophy in human skeletal muscle, particularly in the context of concurrent training.

To circumvent the potentially confounding influence of training status on the mode-specificity of post-exercise molecular responses in skeletal muscle ^21^,22, we investigated potential interference to mTORC1 signalling following exercise protocols that participants were accustomed to via eight weeks of prior training. In contrast to previous studies in untrained or relatively training-unaccustomed participants 14,16-18, we observed enhanced mTORC1 signalling after resistance training compared with concurrent exercise, including greater mTOR and p70S6K1 phosphorylation at 1 h post-exercise, and rps6 phosphorylation at 3 h post exercise. These observations contrast with previous data, including our own ^20^, showing no differences in mTORC1 signalling responses to single bouts of resistance exercise, performed alone or after a bout of continuous endurance exercise ^13^. It has been suggested that any tendency for mTORC1 signalling responses (e.g., p70S6K^Thr389^ phosphorylation) to be further enhanced by concurrent exercise (relative to resistance exercise alone) before training, as shown in a previous study ^14^, were lessened when exercise was performed in a training accustomed state ^13^. While the observed mTORC1 signalling responses were consistent with the paradigm of enhanced mode-specificity of molecular responses with repeated training, the finding of greater AMPK phosphorylation following resistance exercise compared with concurrent exercise was unexpected, given the energy-sensing nature of AMPK signalling and its role in promoting an oxidative skeletal muscle phenotype 48. This observation may suggest an adaptive response whereby endurance training rendered subjects in the concurrent training groups less susceptible to exercise-induced metabolic perturbation in skeletal muscle, manifesting in an attenuated post-exercise AMPK phosphorylation response. A similar phenomenon has been observed in human skeletal muscle after only 10 days of endurance training, whereby post-exercise Taken together, these data lend support to the notion the molecular signals initiated by exercise in skeletal muscle become more mode-specific with repeated training, and increases in post-exercise mTORC1 signalling with concurrent exercise may be attenuated when performed in a training-accustomed state.

While the observed mTORC1 signalling responses were consistent with the paradigm of enhanced mode-specificity of molecular responses with repeated training, the finding of greater AMPK phosphorylation following resistance exercise compared with concurrent exercise was unexpected, given the energy-sensing nature of AMPK signalling and its role in promoting an oxidative skeletal muscle phenotype ^48^. This observation may suggest an adaptive response whereby endurance training rendered subjects in the concurrent training groups less susceptible to exercise-induced metabolic perturbation in skeletal muscle, manifesting in an attenuated post-exercise AMPK phosphorylation response. A similar phenomenon has been observed in human skeletal muscle after only 10 days of endurance training, whereby post-exercise increases in AMPK activity following a single pre-training exercise bout are attenuated compared with the same exercise bout performed before training ^49^. It should also be acknowledged that while AMPK Thr^172^ phosphorylation alone does not necessarily reflect changes in AMPK activity *per se*, ACC Ser^79^ phosphorylation is generally accepted as a marker for AMPK activity ^50^,51. Since we observed greater increases in ACC Ser^79^ phosphorylation with concurrent exercise versus resistance exercise alone during the post-training exercise trial, this may instead reflect further increases in AMPK activity in response to concurrent exercise. Nevertheless, the present data suggest further work is required to define the mode-specificity of AMPK signalling in skeletal muscle and the effect of repeated training on these responses.

In addition to mediating transient changes in translational efficiency, accumulating evidence suggests mTORC1 also plays a key role in regulating ribosome biogenesis (and therefore translational capacity) in skeletal muscle by regulating all three classes of RNA polymerases (RNA Pol-I to -III) ^25^. In agreement with mTORC1 signalling responses, the phosphorylation of upstream regulators of RNA Pol-I-mediated rDNA transcription, including UBF and TIF 1A, was increased more by resistance exercise alone than when combined with endurance exercise in the form of either HIT or MICT. Previous work has demonstrated single sessions of resistance exercise to induce robust increases in TIF-1A Ser^649^ phosphorylation and UBF protein content in human skeletal muscle at 1 h post-exercise, both in untrained and trained states ^34^. Moreover, whereas a single session of resistance exercise did not influence UBF Ser^388^ phosphorylation, this response was elevated in the basal state post-training ^34^. The present data add to the growing body of evidence that resistance exercise is a potent stimulus for increasing the phosphorylation of regulators of Pol-I-mediated rDNA transcription, and suggest these early signalling responses may be similarly attenuated when resistance exercise is combined with endurance exercise. These responses also indicate an apparent disconnect between the upstream signalling responses in the post-training exercise trial related to 45S pre-rRNA transcription (i.e., TIF-1A and UBF phosphorylation), and the basal training-induced changes in markers of ribosomal content (i.e., total RNA and expression of mature rRNA species).

While these responses appear paradoxical, they may suggest that although short-term concurrent training may optimise ribosome biogenesis adaptation versus resistance training performed alone, ribosome biogenesis may instead be further enhanced by longer-term resistance training performed alone. This notion aligns with recent discussion regarding the progression of adaptation with concurrent versus single-mode training, suggesting early adaptation to combined resistance and endurance training may initially be complimentary, whereas longer-term training exacerbates interference to hallmark resistance training adaptations ^52^. Clearly, longer-term training studies are likely required to fully elucidate the effect of concurrent training versus resistance training alone on ribosome biogenesis adaptation in skeletal muscle.

Despite the present findings regarding signalling responses upstream of 45S pre-rRNA transcription, the expression of 45S pre-RNA, but not mature ribosome species, was increased only after concurrent exercise during the post-training exercise trial. Previous work in humans has reported basal increases in 45S pre-rRNA after 8 weeks of resistance training ^34^, and 4 h after a single session of resistance exercise performed in both untrained and trained states ^33^. Notably, post-exercise expression of 45S pre-rRNA was less pronounced in the trained compared with untrained state ^33^. While no substantial basal changes in 45S pre-rRNA expression were observed in the present study, the change in 45S pre-rRNA levels between PRE-T and POST-T was greater for both concurrent training groups compared with RT performed alone. Concurrent exercise also increased 45S pre-rRNA levels at 3 h post-exercise, with little effect of single-mode resistance exercise. These observations may be explained by the muscle sampling time points employed in the present study. Increased post-exercise 45S pre-rRNA levels have been previously shown 4 h after resistance exercise ^33^, whereas a reduction in 45S rRNA levels has been demonstrated 1 h post-resistance exercise in trained, but not untrained, states ^34^. The possibility therefore exists that resistance exercise may increase 45S rRNA expression at a later timepoint post-exercise, and the sampling time points employed herein were not extensive enough to measure any exercise-induced increases in 45S pre-rRNA expression.

The regulation of several Pol-I associated proteins was also measured at the transcriptional level, including TIF-1A, POLR1B, UBF, and cyclin D1. Concurrent exercise, irrespective of endurance training intensity, was sufficient to increase POLR1B mRNA expression at 3 h post exercise, but only MICT+RT and RT alone increased TIF-IA mRNA content at this timepoint. Previous work in human skeletal muscle has demonstrated no effect of a single session of resistance exercise performed in either untrained or trained states on the mRNA expression of either TIF-1A or POLR1B at either 1 h ^34^ or 4 h ^33^ post-exercise. Eight weeks of resistance training has previously been shown to increase basal UBF mRNA expression, which was reduced 1 h following a single session of resistance exercise performed post-training ^34^. Although we observed no basal training-induced increases in UBF mRNA expression for any training group, a similar reduction in UBF mRNA content was noted 3 h post-exercise for the RT group. Increased cyclin D1 mRNA was also seen at rest post-training for the HIT+RT group, which was maintained at 3 h post-exercise. Figueiredo et al. ^34^ have shown eight weeks of resistance training decreased post-training levels of cyclin D1 mRNA compared with pre training, with a small increase induced at 1 h post-exercise by a single session of post-training resistance exercise. It therefore appears HIT is a more potent stimulus for increasing levels of cyclin D1 mRNA compared with resistance exercise alone or MICT, although an acute reduction in cyclin D1 protein levels was also seen 1 h following a single bout of HIT+RT. Previous work has shown increases in cyclin D1 mRNA during long-term (3 months) resistance training ^53^, which may suggest an increase in satellite cell activation and proliferation during the training intervention ^53^,54, although direct measures of these markers were not made in the present study.

The rRNA primers used in the present study were specifically designed to differentiate between mature rRNA expression and the expression of these sequences when still bound to the polycistrionic 45S rRNA precursor (i.e., 5.8S, 18S and 28S [span] rRNA) ^34^. Previous work using these primer sequences has shown basal training-induced increases in mature rRNA expression did not occur concomitantly with increased expression of rRNA transcripts still bound to the 45S precursor (i.e., 5.8S, 18S and 28S [span]), suggesting a training-induced increase in mature rRNA content, rather than increased 45S precursor expression ^34^. In contrast, we observed simultaneous post-exercise increases in the expression of both mature rRNA transcripts and those still bound to the 45S precursor (i.e., ‘span’ rRNA transcripts). It is therefore possible our observed changes in these markers may be reflective solely of changes in 45S pre-rRNA content, and not the mature forms of these rRNAs. However, it is also possible this may relate to the post-exercise time course examined in the present study. In support of this notion, it was shown that a single session of resistance exercise was sufficient to increase only the expression of rRNA transcripts still bound to the 45S pre-rRNA, and not mature rRNA species, even after 48 h of post-exercise recovery ^55^. It is therefore plausible that the post-exercise time courses examined in the present study were not extensive enough to measure early post-exercise changes in mature rRNA expression. Clearly, further work is required to investigate the time course of rRNA regulation with training in human skeletal muscle.

Although we have investigated various upstream regulators of 45S pre-rRNA transcription, it is possible other factors may have been differentially regulated by concurrent versus single mode resistance training and may have contributed to the observed changes in ribosome biogenesis markers. For example, CAD (carbamoyl-phosphate synthetase 2, aspartate transcarbamoylase, dihydroorotase) is directly phosphorylated by p70S6K1 and controls the first three steps in *de novo* pyrimidine synthesis ^56^, a necessary process for accommodating the increased demand for RNA and DNA synthesis to support cellular growth. To our knowledge, the regulation of CAD has, however, not been investigated in the context of training-induced skeletal muscle hypertrophy in humans. Future studies should also consider the potential role of CAD in the regulation of skeletal muscle growth in response to resistance and/or concurrent training.

### Conclusions

This is the first study to simultaneously investigate markers of ribosome biogenesis and mTORC1 signalling in human skeletal muscle following concurrent training compared with single-mode resistance training. Contrary to our hypotheses, and recent work in humans ^33,34^, we noted little evidence of ribosome biogenesis in skeletal muscle following eight weeks of resistance training. Rather, training-induced increases in markers of ribosome biogenesis, tended to be greater following concurrent training and were independent of the endurance training intensity employed. This occurred despite a single session of resistance exercise, when performed post-training, being a more potent stimulus for both mTORC1 signalling and phosphorylation of upstream regulators of RNA Pol-1-mediated rDNA transcription (i.e., TIF 1A and UBF). An apparent disconnect was noted between training-induced changes in muscle fibre CSA, of which the small increase in type I fibre CSA induced by resistance training was attenuated when combined with HIT, and changes in total skeletal muscle RNA content. Overall, the present data suggest single-mode resistance exercise performed in a training accustomed state preferentially induces mTORC1 and ribosome biogenesis-related signalling in skeletal muscle compared with concurrent exercise; however, this is not associated with basal post-training increases in markers of ribosome biogenesis. The observation that both mTORC1 and ribosome biogenesis-related signalling were impaired in response to the final training session of the study for both forms of concurrent exercise, relative to resistance exercise performed alone, suggests resistance training may be a more potent stimulus for ribosome biogenesis and muscle hypertrophy if training were continued longer-term. Further work in human exercise models that stimulate more robust skeletal muscle hypertrophy (e.g., high-volume resistance training performed to failure), together with longer training periods, are likely needed to definitively elucidate the role of ribosome biogenesis in adaptation to resistance training, and subsequently any potential interference to these responses with concurrent training.

## 5. Methods

### Ethical approval

All study procedures were approved by the Victoria University Human Research Ethics Committee (HRE 13-309). After being fully informed of study procedures and screening for possible exclusion criteria, participants provided written informed consent. All methods were performed in accordance with the relevant guidelines and regulations of the Victoria University Human Research Ethics Committee.

### Experimental overview

Participant details and procedures performed in this study have been previously described ^44^; however, these are briefly summarised as follows. The study employed a repeated-measures, parallel-group design (Figure 8A). After preliminary testing for maximal (1-RM) strength, aerobic fitness (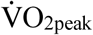, the lactate threshold [LT] and peak aerobic power [Wpeak]), and body composition (dual-energy x-ray absorptiometry [DXA]), participants were ranked by baseline 1-RM leg press strength and randomly allocated to one of three training groups. Each group performed training sessions that consisted of either 1) high-intensity interval training (HIT) cycling combined with resistance training (HIT+RT group, *n* = 8), 2) moderate-intensity continuous training (MICT) cycling combined with resistance training (MICT+RT group, *n* = 7) or 3) resistance training performed alone (RT group, *n* = 8).

**Figure 8.**
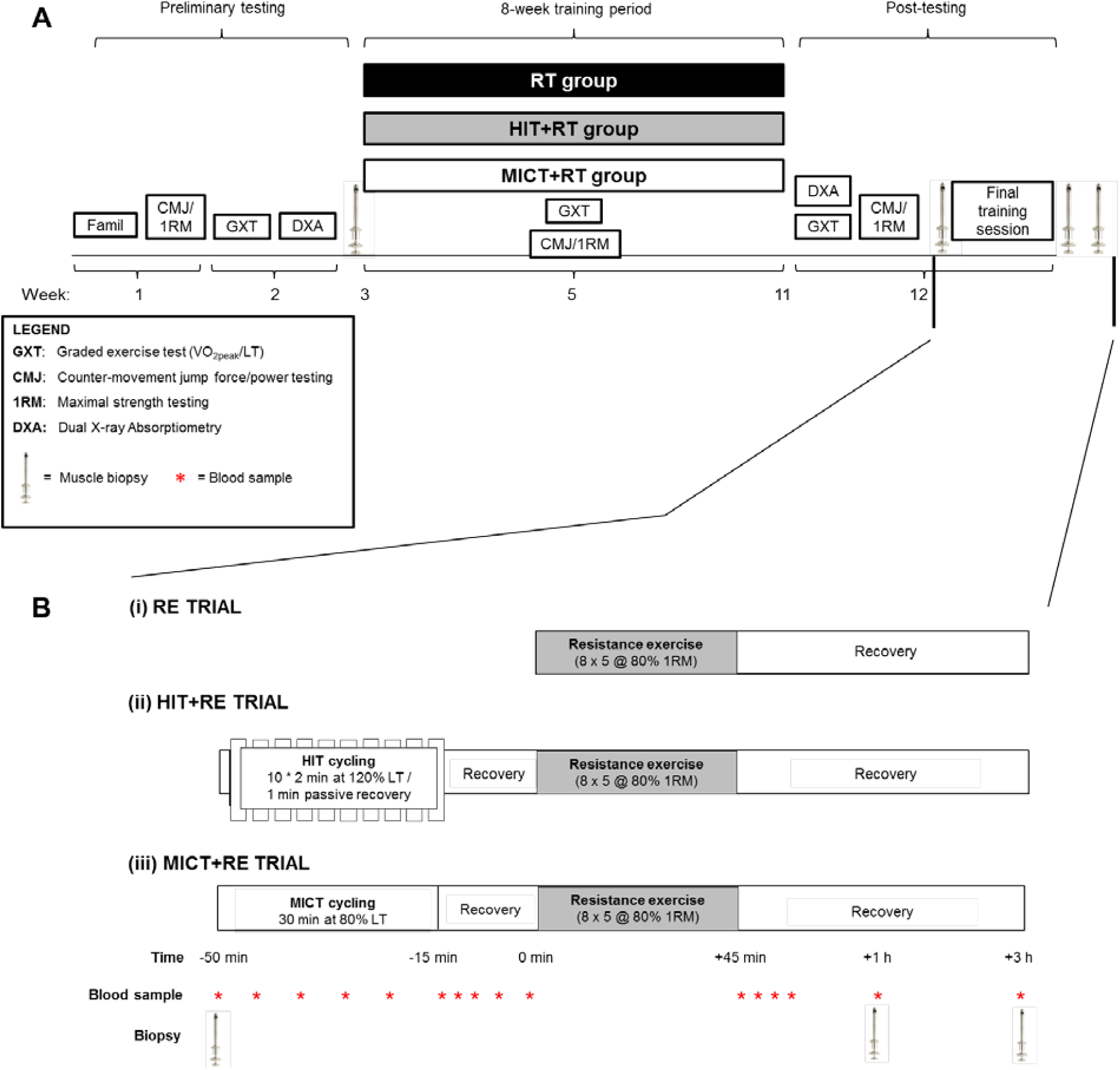
Study overview (A) and timelines for the final training session (B). Participants first completed 8 weeks of either resistance training (RT) alone, or RT combined with either high-intensity interval training (HIT+RT) or moderate-intensity continuous training (MICT+RT). For the final training session (B), participants completed the RE protocol alone (i) or after a 15-min recovery following the completion of either HIT (ii) or work-matched MICT (iii) cycling. Muscle biopsies were obtained from the vastus lateralis at rest before training, and immediately before beginning the final training session, and 1 h and 3 h after completion of RE.

After preliminary testing, and immediately prior to the first training session (i.e., at least 72 h after completion of preliminary testing), a resting muscle biopsy (PRE-T) was obtained from the *vastus lateralis* using the percutaneous needle biopsy technique ^57^ modified with suction ^58^. Participants then completed 8 weeks of group-specific training performed three times per week. Between 48 and 72 h after completing the post-training 1-RM strength testing, participants underwent a final group-specific training session (Figure 8B) whereby early post-exercise molecular responses in skeletal muscle were measured in a training-accustomed state. Three additional biopsies [at rest (POST-T), and 1 h (+1 h) and 3 h (+3 h) post-exercise] were obtained during the final group-specific training session.

### Training intervention

The training intervention in this study has previously been described in detail ^44^. Briefly, participants began the 8-week training intervention 3 to 5 days after completion of preliminary testing. All training groups performed an identical resistance training program on non consecutive days (typically Monday, Wednesday, and Friday), with the HIT+RT and MICT+RT groups completing the corresponding form of endurance exercise 10 min prior to commencing each resistance training session.

### Final training session

Two or three days after completion of the training intervention and post-testing, participants perfomed a final group-specific training session (Figure 8B) whereby early post-exercise skeletal muscle responses were measured in a training-accustomed state. Participants reported to the laboratory after an overnight (∼ 8-10 h) fast. After resting quietly for ∼ 15 min upon arrival at the laboratory, a venous cathether was inserted into an anticubital forearm vein and a resting blood sample was obtained. A resting, post-training (POST-T) muscle biopsy was then taken from the *vastus lateralis* muscle (described subsequently). Participants in the RT group waited quietly for 10 min after the POST-T biopsy and then completed a standardised resistance exercise protocol (8 x 5 leg press repetitions at 80% of the post-training 1-RM, three minutes of recovery between sets). Participants in the HIT+RT and MICT+RT groups preceded the standardised RT with either HIT (10 x 2-min intervals at 140% of the post-training LT, 1 min passive recovery between intervals) or work- and duration-matched MICT cycling (30 min at 93.3% post-training LT), respectively. Fifteen minutes of passive recovery was allowed between completion of either HIT or MICT and the subsequent resistance exercise bout. Each cycling bout was performed after a standardised warm-up ride at 75 W for 5 min. After completion of resistance exercise, participants rested quietly in the laboratory and additional biopsies were obtained after 1 (+1 h) and 3 h (+3 h) of recovery. Venous blood samples were also obtained at regular intervals during cycling and following recovery from both cycling and resistance exercise (Figure 8B).

### Muscle sampling

After administration of local anaesthesia (1% Xylocaine), a small incision (∼ 7 mm in length) was made through the skin, subcutaneous tissue, and fascia overlying the *vastus lateralis* muscle for each subsequent biopsy. A 5-mm Bergström needle was then inserted into the muscle and a small portion of muscle tissue (∼ 50-400 mg) removed. All biopsies were obtained from separate incision sites in a distal-to-proximal fashion on the same leg as the pre-training biopsy. Muscle samples were blotted on filter paper to remove excess blood, immediately frozen in liquid nitrogen, and stored at −80°C until subsequent analysis. A small portion of each biopsy sample (∼ 20 mg) was embedded in Tissue-Tek (Sakura, Finetek, NL), frozen in liquid nitrogen-cooled isopentane, and stored at −80°C for subsequent immunofluorescence analysis.

### Western blotting

Approximately 5 mg of frozen muscle tissue was homogenised in lysis buffer (0.125M Tris-HCl, 4% SDS, 10% Glycerol, 10mM EGTA, 0.1M DTT, 1% protease/phosphatase inhibitor cocktail), left for 1 h at room temperature, and then stored overnight at −80°C. The following morning, samples were thawed and the protein concentration determined (Red 660 Protein Assay Kit, G-Biosciences, St. Louis, MO). Bromophenol blue (0.1%) was then added to each sample, which were then stored at −80°C until subsequent analysis. Proteins (8 μg) were separated by SDS-PAGE using 6-12% acrylamide pre-cast gels (TGX Stain Free, Bio-Rad laboratories, Hercules, CA) in 1× running buffer (25 mM Tris, 192 mM Glycine, 0.1% SDS), and transferred to polyvinylidine fluoride (PVDF) membranes (Bio-Rad laboratories, Hercules, CA) using a semi-dry transfer system (Trans Blot Turbo, Bio-Rad laboratories, Hercules, CA) for 7 min at 25 V. After transfer, membranes were blocked with 5% skim milk in 1×TBST (200 mM Tris, 1.5 M NaCl, 0.05% Tween 20) for 1 h at room temperature, washed with 1×TBST (5×5 min), and incubated with primary antibody solution (5% BSA [bovine serum albumin], 0.05% Na Azide in 1×TBST) overnight at 4°C. Primary antibodies for phosphorylated (p-) p mTOR^Ser2448^ (1:1000; #5536), mTOR (1:1000), p-p70S6K1^Thr389^ (1:1000; #9234), p70S6K1 (1:1000), p-4E-BP1^Thr37/46^ (1:1000; #2855), 4E-BP1 (1:1000; #9452), p-AMPK^Thr172^ (1:1000; #2535), AMPK (1:1000; #2532), p-rps6^Ser235/236^ (1:750; #4856), rps6 (1:1000; #2217) and p ACC^Ser79^ (1:1000; #3661) were from Cell Signalling Technology (Danvers, MA), p-UBF^Ser388^ (1:1000; sc-21637-R), UBF (1:000; sc-9131) and cyclin D1 (1:1000; sc-450) were from Santa Cruz Biotechnology (Dallas, TX), and p-RRN3 (TIF-1A)^Ser649^ (1:1000; ab138651) and TIF-1A (1:1000; ab70560) were from Abcam (Cambridge, UK). The following morning, membranes were washed again with 1×TBST and incubated with a secondary antibody (Perkin Elmer, Waltham, MA, #NEF812001EA; 1:50000 or 1:100000 in 5% skim milk and 1×TBST) for 1 h at room temperature. After washing again with 1×TBST, proteins were detected with chemiluminescence (SuperSignal™ West Femto Maximum Sensitivity Substrate, Thermo Fisher Scientific, Waltham, MA) and quantified via densitometry (Image Lab 5.0, Bio-Rad laboratories, Hercules, CA). Representative western blot images for each protein target analysed are shown in Figure 9. All sample timepoints for each participant were run on the same gel and normalised to both an internal pooled sample present on each gel, and the total protein content of each lane using a stain-free imaging system (Chemi Doc™ MP, Bio-Rad laboratories, Hercules, CA). Phosphorylated proteins were then expressed relative to the total amount of each respective protein (with the exception of phosphorylated ACC^Ser79^, which was normalised only to the total protein content of each lane due to technical difficulties when measuring total ACC protein).

**Figure 9.**
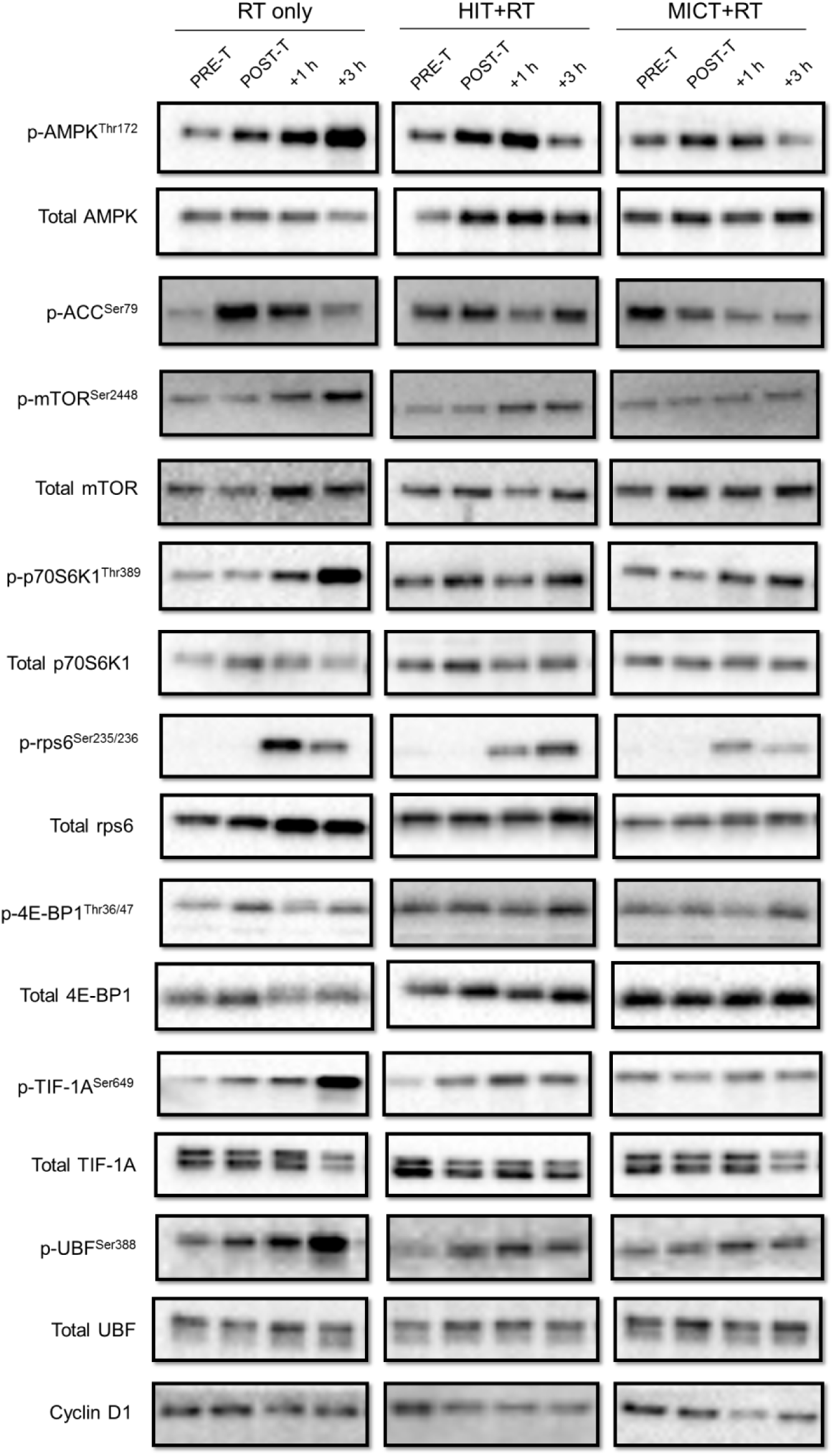
Representative western blots for the phosphorylation (p-) and total protein content of signalling proteins before (PRE-T) and after (POST-T) eight weeks of either RT alone, or RT combined with either high-intensity interval training (HIT+RT) or moderate-intensity continuous training (MICT+RT), and 1 h (+1 h) and 3 h (+3 h) after a single exercise bout performed post-training. Cropped western blot images are displayed for clarity of presentation, and full-length western blot images are presented in supplementary information.

### Real-time quantitative PCR (qPCR)

#### RNA extraction

Total RNA (1145 ± 740 ng; mean ± SD) was extracted from approximately 25 mg of muscle tissue using TRI Reagent® (Sigma Aldrich, St. Louis, MO) according to the manufacturer’s protocol. Muscle samples were firstly homogenised in 500 μL of TRI Reagent® using a Tissue Lyser II and 5 mm stainless steel beads (Qiagen, Venlo, Limburg, Netherlands) for 120 s at 30 Hz. After resting for 5 min on ice, 50 μL of 1-bromo-3-chloropropane (BCP) was added to the tube, inverted for 30 s to mix, and then rested for 10 min at room temperature. The homogenate was then centrifuged for 15 min at 13,000 rpm and the upper transparent phase transferred to another tube. Isopropanol (400 μL) was added to the tube, inverted briefly to mix, and stored overnight at −20°C to precipitate the RNA. After overnight incubation, the solution was centrifuged for 60 min at 13,000 rpm and at 4°C to pellet the RNA. The RNA pellet was washed twice by centrifugation in 75% ethanol/nuclease-free water (NFW) for 15 min at 13,000 rpm, allowed to air-dry, and then dissolved in 15 μL of NFW (Ambion Inc., Austin, TX). The quantity and quality of RNA was subsequently determined using a spectrophotometer (NanoDrop One, Thermo Scientific, Wilmington, DE). The purity of RNA was assessed using the ratio between the absorbance at 260 nm and absorbance at 280 nm (mean ± SD; 2.37 ± 0.43), and the ratio between the absorbance at 260 nm and absorbance at 230 nm (1.71 ± 0.42). The total skeletal muscle RNA concentration was calculated based on the total RNA yield relative to the wet weight of the muscle sample.

### Reverse transcription

For mRNA analysis, first-strand cDNA was generated from 1 μg RNA in 20 μL reaction buffer using the iScript® cDNA synthesis kit (Bio-Rad laboratories, Hercules, CA) according to manufacturer’s protocol, with each reaction comprising 4 μL 5× iScript reaction mix, 1 μL iScript Reverse Transcriptase, 5 μL NFW and 10 μL of RNA sample (100 ng/μL). Reverse transcription was then performed with the following conditions: 5 min at 25°C to anneal primers, 30 min at 42°C for the extension phase, and 5 min at 85°C. Following reverse transcription, samples were DNase-treated (Life Technologies, Carlsbad, CA) and cDNA was stored at −20°C until further analysis.

### Real-time quantitative PCR (qPCR)

Real-time PCR was performed using a Realplex^2^ PCR system (Eppendorf, Hamburg, Germany) to measure mRNA levels of UBF, TIF-1A, cyclin D1, POLR1B, and commonly used reference genes GAPDH (glyceraldehyde 3-phosphate dehydrogenase), cyclophilin (also known as peptidyl-prolylcis-trans isomerase), β2M (beta-2 microglobulin) and TBP (TATA binding protein). Target rRNAs were the mature ribosome species 5.8S, 18S and 28S. Since primers specific for these mature rRNA sequences will also amplify pre-RNA transcripts (i.e., the 45S pre-rRNA), we used specifically designed primers (QIAGEN, Venlo, Limburg, The Netherlands) to distinguish between mature rRNA species and those still bound to the 45S pre rRNA transcript, as previously described ^34^. Briefly, primers were designed specifically for pre-rRNA sequences spanning the 5’end external/internal transcribed spacer regions (ETS and ITS, respectively) of the 45S pre-RNA transcript and the internal regions of mature rRNA sequences (i.e., 18S-ETS, 5.8S-ITS, and 28S-ETS). For clarity, primers amplifying the mature rRNA transcripts are henceforth designated as ‘mature’ transcripts (e.g., 18S rRNA [mature]), as opposed to those primers amplifying rRNA sequences bound to the 45S rRNA precursor, henceforth designated as ‘span’ transcripts (e.g., 18S rRNA [span]). A specific primer for the initial region of the 5’ end of the 45S pre-rRNA transcript was used to measure 45S pre-rRNA expression levels ^34^. Standard and melting curves were performed for all primers to ensure both single-product and amplification efficiency. Details for all primers used are provided in Table 4 (mRNA) and Table 5 (rRNA).

**Table 4.**
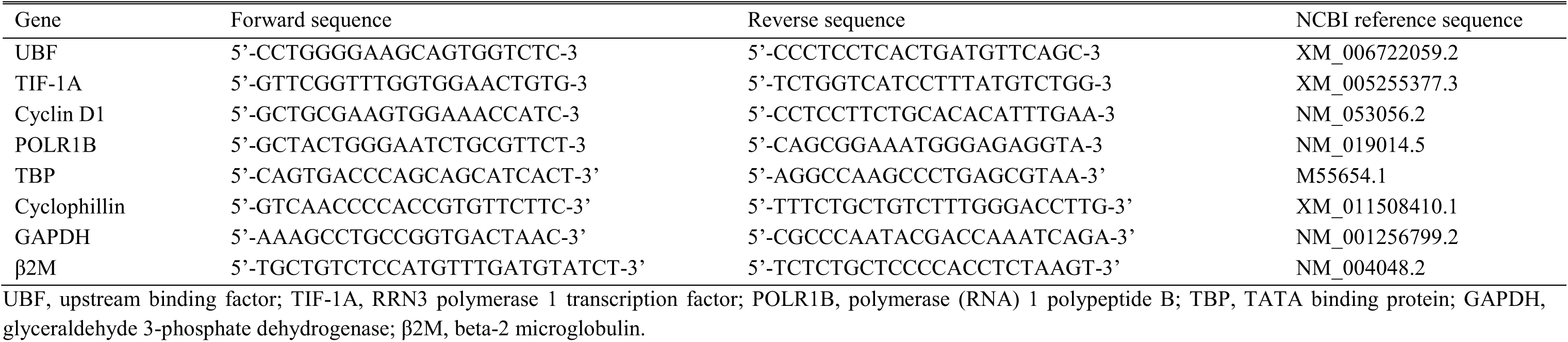
Details of PCR primers used for mRNA analysis

**Table 5.** Details of PCR primers used for rRNA analysis

| Target | Catalogue number |
| --- | --- |
| 45S pre-rRNA | PPH82089A |
| 5.8S rRNA (mature) | PPH82091A |
| 18S rRNA (mature) | PPH71602A |
| 28S rRNA (mature) | PPH82090A |
| 5.8S-ITS (span) | PPH82111A |
| 18S-ETS (span) | PPH82110A |
| 28S-ITS (span) | PPH82112A |

Each PCR reaction was performed in duplicate using a robotic pipetting machine (EpMotion 2100, Eppendorf, Hamburg, Germany) in a final reaction volume of 10 μL containing 5.0 μL 2× SYBR green (Bio-Rad Laboratories, Hercules, CA), 0.6 μL PCR primers (diluted to 15 μM; Sigma Aldrich, St. Louis, MO), 0.4 μL NFW and 4 μL cDNA sample (diluted to 5 ng/μL). Conditions for the PCR reactions were: 3 min at 95°C, 40 cycles of 15 sec at 95°C/1 min at 60°C, one cycle of 15 sec at 95°C/15 sec at 60°C, and a ramp for 20 min to 95°C. Each plate was briefly centrifuged before loading into the PCR machine. To compensate for variations in input RNA amounts and efficiency of the reverse transcription, mRNA data were quantified using the 2^−ΔΔCT^ method ^59^ and normalised to the geometric mean ^60^ of the three most stable housekeeping genes analysed (cyclophillin, β2M and TBP), determined as previously described ^61^.

### Immunohistochemistry

Muscle cross-sections (10 μM) were cut at −20°C using a cryostat (Microm HM 550, Thermo Fisher Scientific, Waltham, MA), mounted on uncoated glass slides, and air-dried for 20 min at room temperature. Sections were then rinsed briefly with 1×PBS (0.1M; Sigma Aldrich, St Louis, MO), fixed with cold paraformaldehyde (4% in 1×PBS) for 10 min at room temperature, rinsed three times with 1×PBS, incubated in 0.5% TritonX in 1×PBS for 5 min at room temperature, rinsed again three times with 1×PBS, and then blocked for 1 h at room temperature in a 3% BSA solution in 1×PBS. After blocking, sections were then incubated with a primary antibody for myosin heavy chain type I (A4.840, Developmental Studies Hybridoma Bank, University of Iowa, IA), diluted 1:25 in 3% BSA/PBS overnight at 4°C. The following morning, sections were washed four times in 1×PBS for 10 min each, before incubating with a secondary antibody (Alexa Fluor® 488 conjugate Goat anti-mouse IgM, cat. no. A-21042, Thermo Fisher Scientific, Waltham, MA) diluted 1:200 in 3% BSA/PBS for 2 h at room temperature. Sections were again washed four times in 1×PBS for 10 min each, before incubation with Wheat Germ Agglutinin (WGA) (Alexa Fluor® 594 Conjugate; cat. no. W11262, Thermo Fisher Scientific, Waltham, MA), diluted to 1:100 in 1×PBS (from a 1.25 mg/mL stock solution), for 15 min at room temperature. Sections were washed again 4 times with 1×PBS for 3 min each, blotted dry with a Kim-Wipe, and Flouroshield™ (cat. no. F6182; Sigma Aldrich, St Louis, MO) added to each section before the coverslip was mounted. Stained muscle sections were air-dried for ∼ 2 h and viewed with an Olympus BX51 microscope coupled with an Olympus DP72 camera for flourescence detection (Olympus, Shinjuku, Japan). Images were captured with a 10× objective and analysed using Image Pro Premier software (version 9.1; Media Cybernetics, Rockville, MD). Analysis was completed by an investigator blinded to all groups and time points. For each subject, muscle fibre CSA was determined for both type I and type II muscle fibres. For the RT, HIT+RT and MICT+RT groups, a total of 107 ± 61, 112 ± 67, and 84 ± 73 (mean ± SD) type I fibres and 154 ± 72, 136 ± 80, and 144 ± 76 (mean ± SD) type II fibres were included for analysis, respectively.

### Statistical analyses

The effect of training group on outcomes was analysed using a combination of both traditional and magnitude-based statistical analyses. Western blot, qPCR and immunohistochemistry data were log-transformed before analysis to reduce non-uniformity of error ^62^. Data were firstly analysed via a two-way (time × group) analysis of variance with repeated-measures (RM-ANOVA) (SPSS, Version 21, IBM Corporation, New York, NY). To quantify the magnitude of within- and between-group differences for dependent variables, a magnitude-based approach to inferences using the standardised difference (effect size, ES) was used ^62^. The magnitude of effects were defined according to thresholds suggested by Hopkins ^62^, whereby <0.2 = trivial, 0.2-0.6 = small, 0.6-1.2 = moderate, 1.2-2.0 = large, 2.0-4.0 = very large and >4.0 = extremely large effects. Lacking information on the smallest meaningful effect for changes in protein phosphorylation and gene expression, the threshold for the smallest worthwhile effect was defined as an ES of 0.4, rather than the conventional threshold of 0.2 ^20^. Magnitude-based inferences about effects were made by qualifying the effects with probabilities reflecting the uncertainty in the magnitude of the true effect ^63^. Effects that were deemed substantial in magnitude (and therefore meaningful) were those at least 75% ‘*likely*’ to exceed the smallest worthwhile effect (according to the overlap between the uncertainty in the magnitude of the true effect and the smallest worthwhile change ^63^). Exact *P* values were also determined for each comparison, derived from paired (for within-group comparisons) or unpaired (for between-group comparisons) *t*-tests, with a Bonferroni correction applied to correct for multiple comparisons (SPSS, Version 21, IBM Corporation, New York, NY). A summary of all within- and between-group comparisons for this study are presented in supplementary tables 1 and 2, respectively. Physiological (blood lactate, blood glucose, heart rate) and psychological (rating of perceived exertion [RPE]) responses to exercise are reported as mean values ± SD, whereas protein phosphorylation and gene expression data are reported as mean within- and between-condition percentage differences ±90 % CL.

## Data availability

The datasets generated during and/or analysed during the current study are available from the corresponding author upon reasonable request.

## 7. Acknowledgements

We gratefully acknowledge the efforts of the participants, without whom this study would not have been possible. We also acknowledge Dr Chris Shaw (Deakin University) for technical assistance with the immunofluorescence analysis. This study was supported, in part, by a grant from the Gatorade Sports Science Institute (GSSI) awarded to J.J.F.

## 8. Author contributions statement

Study design was performed by J.J.F., J.D.B., E.D.H., D.J.B. and N.K.S. Data collection was performed by J.J.F, M.J.A and A.P.G. Analysis and interpretation of data was performed by J.J.F., J.D.B., E.D.H., D.J.B. and N.K.S. The manuscript was written by J.J.F., D.J.B., and N.K.S., while J.D.B., E.D.H., M.J.A and A.P.G. critically revised the manuscript. All authors approved the final version of the manuscript.

All data collection and data analysis for this study was conducted and performed in the exercise physiology and biochemistry laboratories at Victoria University, Footscray Park campus, Melbourne Australia.

